# The Molecular Structure of Anterograde Intraflagellar transport trains

**DOI:** 10.1101/2022.08.01.502329

**Authors:** Samuel E. Lacey, Helen E. Foster, Gaia Pigino

**Affiliations:** Human Technopole, Viale Rita Levi-Montalcini, 1, 20157 Milan, Italy

## Abstract

Anterograde intraflagellar transport trains are essential for cilia assembly and maintenance. These trains are formed of 22 IFTA and IFTB proteins that link structural and signalling cargoes to microtubule motors for import into cilia. It remains unknown how the IFTA/B proteins are arranged into complexes and how these complexes polymerise into functional trains. Here, we use *in situ* cryo-electron tomography and Alphafold2 protein structure predictions to generate the first molecular model of the entire anterograde train. We show how the conformation of both IFTA and IFTB is dependent on lateral interactions with neighbouring repeats, suggesting that polymerization is required to cooperatively stabilize the complexes. The retrograde dynein motor binding site is a composite surface involving multiple IFTB repeats, ensuring that dynein can only form a strong interaction with IFTB upon train assembly. Finally, we reveal how IFTB extends two flexible tethers to maintain a connection with IFTA that can withstand the mechanical stresses present in actively beating cilia. Overall, our findings provide a framework for understanding the fundamental processes that are involved in cilia assembly.

## Introduction

Cilia are hair-like organelles that extend from a cell and beat back and forth to create motion (motile cilia) or act as a hub for inter-cell signalling (primary cilia). At their core is a ring of nine interconnected microtubule doublets (MTs) in a well-characterised structure known as the axoneme (**Figure 1A)**. A diffusion barrier exists at the base of the cilium, meaning that the vast quantities of structural proteins required to build the axoneme need to be delivered by microtubule motors in a process called intraflagellar transport (IFT). In primary cilia, IFT also transports membrane-associated proteins into and out of the cilium to regulate key developmental signalling pathways^1^. Underlining the importance of IFT, the absence of many IFT proteins is lethal, and mutations in IFT-related proteins can result in a group of congenital diseases called “ciliopathies”, with diverse developmental phenotypes^2^.

**Figure 1.**
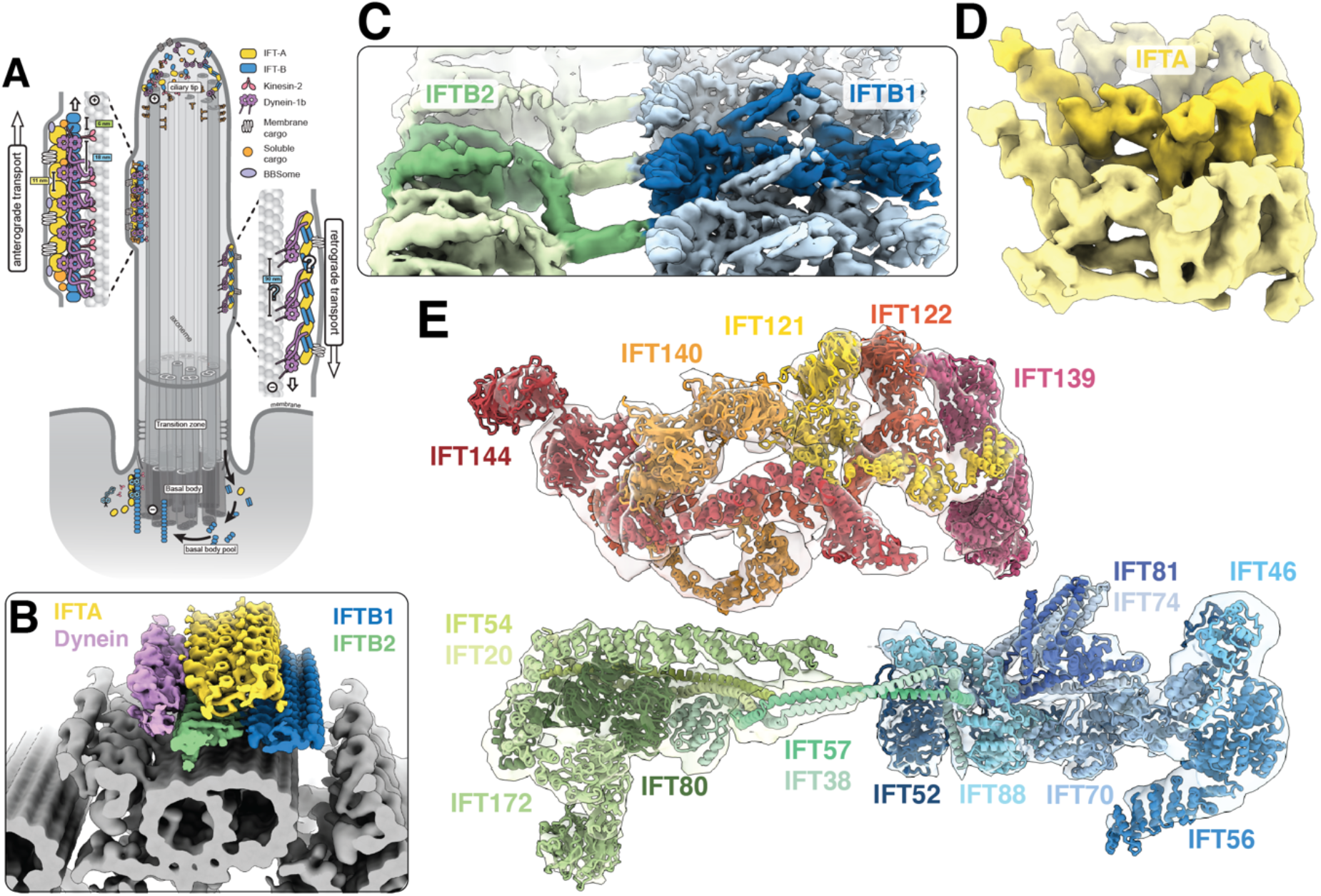
An overview of the anterograde IFT train structure. **A –** Cartoon model of IFT within a cilium. Anterograde trains form at the base of the cilium (basal body) and carry cargo through the diffusion barrier (transition zone) and to the tip. Here, they remodel into retrograde trains that carry their cargoes back to the basal body for recycling. **B** − The new subtomogram averages lowpass filtered and coloured by complex (IFTA yellow, IFTB1 blue, IFTB2 green, dynein purple), docked onto a cryo-ET average of the microtubule doublets found in motile cilia. One repeating unit is highlighted in each complex with darker shading **C –** The new subtomogram averages for IFTB1 (blue) and IFTB2 (green) displayed together as a composite. One repeating unit is highlighted with darker shading **D** −The new subtomogram average of IFTA. **E** − Following flexible fitting we obtain a molecular model for the entire anterograde IFT train, shown here as if looking down the train. Density for four maps is shown; IFTB2 and IFTA, with the main IFTB1 average combined with a masked refinement of the region containing IFT56 (IFTB1 tail, Figure S2A) since this region is more flexible relative to the core.

IFT is organized by the IFTA and IFTB protein complexes. Together these assemble into ordered and repetitive IFT “trains” that link the microtubule motors to hundreds of different IFT cargoes. The IFT process is initiated at the base of the cilium, where IFTB complexes start to polymerise on their own^3^. This nascent train acts as a platform for IFTA polymerisation, and recruits kinesin-2 motors (**Figure 1A)**. Various structural and signalling cargoes then dock to the train, as well as autoinhibited cytoplasmic dynein-2 motors. The kinesin carries the train into the cilium, and the cargoes dissociate at the tip to be incorporated into the axoneme or diffuse in the ciliary membrane^4,5^. The IFTA/B components then remodel into a conformationally distinct retrograde train, which rebinds to the now-active dynein-2 and transports a new selection of cargoes back to the cell body^6–8^.

From our previous cryo-electron tomography (cryo-ET) study of in-situ *Chlamydomonas reinhardtii* cilia, we know the overall appearance of anterograde trains to ∼30Å resolution^9^. IFTB, which contains 16 proteins (IFT172/88/81/80/74/70/57/56/54/52/46/38/27/25/22/20), forms a 6nm repeat with one autoinhibited dynein-2 bound every third repeat **(Figure 1B)**. IFTA, which contains 6 proteins (IFT144/140/139/122/121/43), sits between IFTB and the membrane. It has an 11.5nm repeat, creating a mismatch in periodicity between IFTA and IFTB polymers. However, due to the limited resolution the molecular architecture of IFTA and IFTB remains unknown. Crystal structures of some IFTB proteins have been solved ^10–15^, but they are mostly of small fractions of the overall proteins. Much of our knowledge therefore comes from biochemically mapped interactions between isolated IFTB proteins ^10,11,16^. None of the six IFTA components have been structurally characterised, and there are fewer verified interactions for this complex^16–18^.

As a result, we have a limited understanding of many fundamental mechanisms underlying IFT. To address this, we generated significantly improved (10-18Å) subtomogram averages of *Chlamydomonas* IFT trains, allowing us to build the first complete molecular model of the anterograde train. Here, we present a tour of the IFTA and IFTB complexes within the context of polymerised trains. Together, our results provide insights into the organisation and assembly of IFT trains, how cargoes are bound to the train, and the conversion of anterograde trains into retrograde trains.

### Creating a model of anterograde IFT trains

To generate a molecular model of the anterograde IFT train, we collected 600 cryo-electron tomograms of *Chlamydomonas* cilia. Anterograde IFT trains were readily identifiable for manual picking as repeating filaments between the microtubule doublets and the membrane (**Figure S1**). We picked and refined IFTB and IFTA repeats independently due to their periodicity mismatch, and performed subtomogram averaging with the STOPGAP-Warp/M-Relion3 processing pipeline (**Figure S2-4)**. In IFTB, we identified two rigid bodies that flex around a central hinge that correspond to the biochemically characterized IFTB1 and IFTB2 sub-complexes (**Figure S2A)**. After masked refinements, we obtained structures at 9.9Å resolution for IFTB1, 11.5Å for IFTB2, and 18.6Å for IFTA (**Figure 1C/D, Figure S3A-D, Supplementary table 1)**.

To understand how the IFT proteins are organised in their complexes, we built a molecular model into our maps. As de novo model building is not possible at this resolution, we used a hybrid approach by flexibly fitting high confidence Alphafold2 models of IFT proteins **(Supplementary table 2)**. This allowed us to build a molecular model of the complete anterograde train (**Figure 1E, Figure S5B, Supplemental movie 1-3**).

### IFTB is organised around IFT52

IFTB is central to the assembly of anterograde trains. It recruits active kinesin motors, and carries both the IFTA complex and the retrograde motor dynein-2 to the tip^19^ **(Figure 1B)**. IFTB is also responsible for the recruitment of all characterized structural cargoes, as well as a subset of membrane-bound cargoes, to anterograde trains. It is an elongated complex with two distinct lobes corresponding to the IFTB1 and IFTB2 subcomplexes (**Figure 2A-D**). Our structure reveals the crucial role that the IFTB1 component IFT52 has in the structural integrity of the entire IFTB complex.

**Figure 2.**
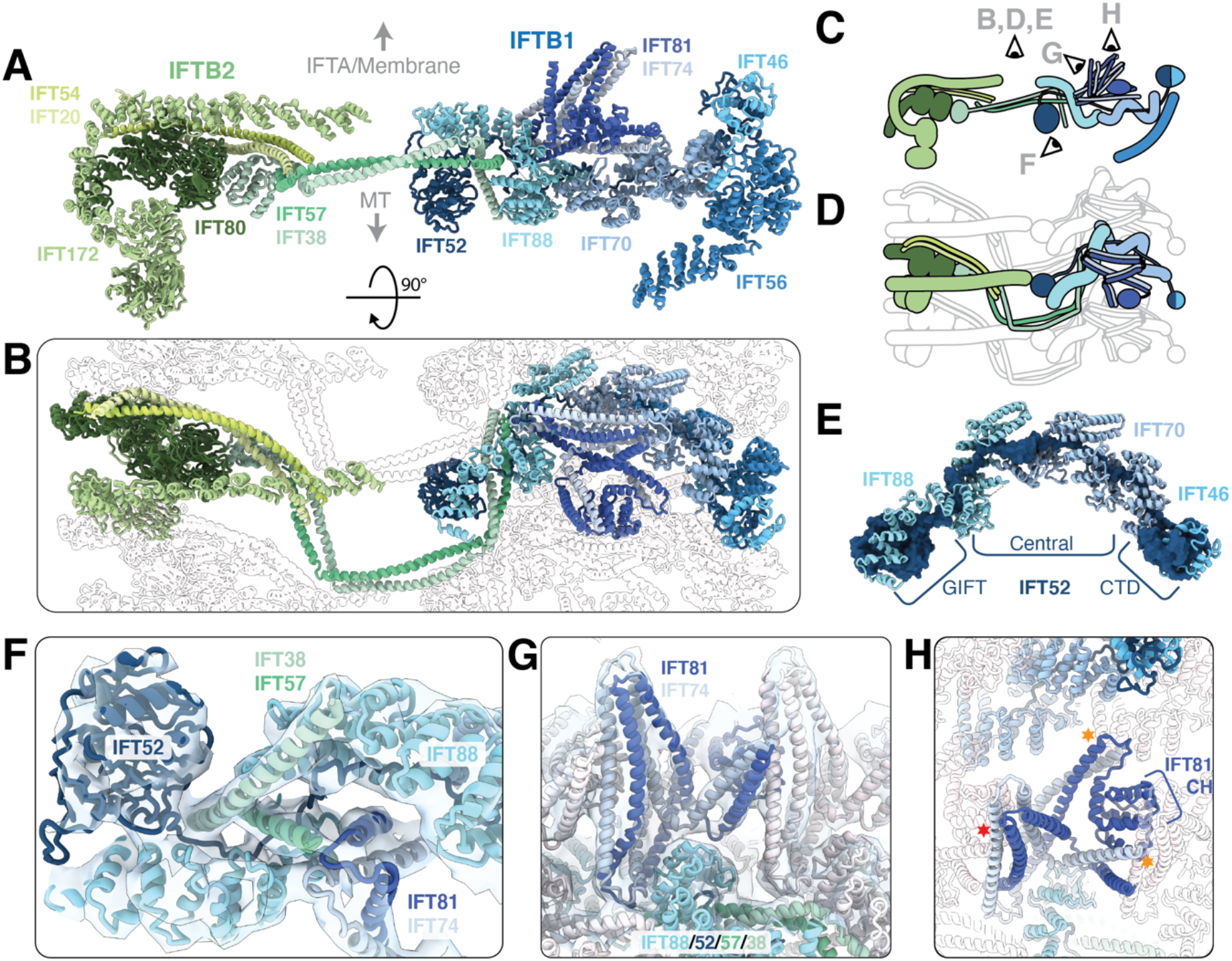
IFT52 is central to the overall IFTB complex. **A** − One repeat of the IFTB complex viewed in profile, looking down the train. **B** − Top view of the IFTB polymer, as if looking down from the membrane/IFTA. A single repeat is shown in colour, with adjacent repeats shown in silhouette. Colouring as in A. **C** − Cartoon representation of **A**, showing the viewing positions of other panels in the figure **D** − Cartoon representation of **B**. **E** − IFT52 (dark blue), shown as a molecular surface, forms the core of the IFTB1 complex, with the central unstructured domain threading through the TPR superhelices of IFT88 (cyan) and IFT70 (steel blue). **F** − IFT57/38 (dark/light green) from IFTB2 interact with IFTB1 by fitting into a cleft in the TPR superhelix of IFT88 (cyan) along with the unstructured IFT52 central domain (dark blue). **G** − IFT81/74 (navy blue/grey) sit on top of IFT88, and form a compressed segmented coiled coil repeating along the IFT train. **H** − Top view of **G**. Lateral interactions to IFT81/74 in adjacent repeats is highlighted with stars (red star to IFT81-CH on N-1 repeat, orange star to IFT81/74-CC and IFT70 of N+1 repeat).

IFT52 consists of an N-terminal GIFT domain, a central disordered region, and a C-terminal domain (CTD) that forms a heterodimer with IFT46^11^ (**Figude 2E, S5A)**. It spans the length of IFTB1, with the GIFT domain on the MT-proximal surface at the center of the train, and the IFT52-CTD:IFT46 heterodimer at the periphery (**Figure 2A/B)**. IFT88 and IFT70, two supercoiled TPR proteins, wrap around the central disordered domain of IFT52 by stacking end-to-end to create a continuous central pore **(Figure 2E, Figure S6A/B/F)**. IFT70 is known to make a tight spiral with a hydrophobic core, and IFT52 is thought to be an integral part of its internal structure^11^. However, we see that IFT88 forms a more open spiral with charged internal surfaces, suggesting that its interaction with IFT52 is reversible. The remainder of IFTB1 is assembled around the IFT88/70/52 trimer, which binds to the coiled coil IFT81/74 subcomplex and IFT56, a third TPR spiral protein (**Figure S6D/E)**. Therefore, the IFTB1 subcomplex is assembled around IFT52.

Additionally, IFT52 and IFT88 form the main interface between IFTB1 and IFTB2. This is mediated through interactions with the IFT57/38 complex of IFTB2, consistent with biochemical data^10^. IFT57/38 is a segmented coiled coil, with both proteins also containing an N-terminal Calponin homology (CH) domain. IFT38-CH was previously shown to form a high affinity interaction with the N-terminal WD domain of IFT80^15^. In our structure this interaction anchors IFT57/38 in IFTB2 (**Figure S6G)**, and the coiled coils then extend across the central region to contact IFT88 from the neighbouring repeat (**Figure 2B)**. Here, a conserved proline residue in both IFT57 and IFT38 creates a right-angled kink (**Figure S6H)** which points the subsequent coiled coil segment towards the IFT88 in the same repeat. The loose spiral of IFT88 creates an open cleft which IFT58/37 and the IFT52 disordered region slots into, creating multiple contacts between the IFTB1 and IFTB2 components **(Figure 2F)**.

Taken together, we find that IFT52 is the cornerstone of the IFTB complex. This is consistent with results from the *Chlamydomonas bld1* mutant, which lacks functional IFT52 and cannot grow cilia or form IFTB complexes as a result^20,21^. Furthermore, in humans a mutation in IFT52 at the interface with IFT57/38 (D259H, corresponding to D268 in *Chlamydomonas* (**Figure S6I)** is associated with a developmental kidney ciliopathy^22^, which could be caused by destabilization in the association of IFTB1 and B2.

### IFT81/74 is stabilized by neighbouring repeats

Next, we wanted to understand how the individual IFTB1 complexes associate as polymers. Part of the interaction is mediated by simple wall-to-wall contacts between adjacent IFT88/70/52 trimers **(Figure 2B)**. These contacts are supplemented by a more intricate network of lateral interactions in the IFT81/74 dimer that sits on top of IFT88/70/52. IFT81/74 forms eight coiled coil segments (CC1-8), and is the binding site for IFT27, IFT25 and IFT22^11,13^. The loop between IFT81/74 CC1 and CC2 forms the main attachment to the IFTB1 core by binding to the same cleft in IFT88 as IFT57/38 **(Figure 2F/G)**. The first four coiled coil segments then form two interactions with adjacent IFTB1 repeats, forcing them into a folded/compressed conformation (**Figure 2H**). First, the N-terminal IFT81-CH domain is raised above the IFT88/70/52 trimer through an interaction between IFT81/74-CC1 and IFT70 of the neighbouring repeat. Then, IFT81-CH acts as a strut against which CC2/3 from the neighbouring repeat leans in an upright position. Since the coiled coil segments are linked by flexible loops, this suggests that a feature of IFTB polymerisation is the cooperative stabilisation of IFT81/74 in a compressed conformation

### Binding sites for IFT27/25/22 are oriented towards the membrane

To complete our understanding of IFTB1, we considered the position of the remaining IFT27, IFT25 and IFT22 subunits. The binding sites for these proteins are on the CC5 to CC8 segments of IFT81/74^11,13^. However, only the first four segments are present in our density, indicating that IFT27/25/22 are flexibly tethered to the IFTB1 polymer. The position of IFT81/74-CC4, the last resolved segment, projects the flexible regions out towards the membrane **(Figure 2A/G)**. This allows IFT27/25/22 to fulfill proposed roles in recruitment of membrane cargoes ^23,24^, and provides sufficient flexibility to maintain an interaction with proteins in the crowded ciliary membrane.

### IFT80 forms the core of IFTB2

The IFTB2 subcomplex forms the second lobe of IFTB (**Figure S7A-D, Supplementary movie 2)**. It is made up of two pairs of coiled coil proteins (IFT57/38 and IFT54/20) and two large proteins (IFT172 and IFT80) which each contain a pair of tandem WD domains followed by C-terminal TPR motifs (**Figure S5A/B**). The second WD domain of both these proteins forms an incomplete circle (**Figure 3A-C**, **Figure S7F)**, particularly dramatically in the case of IFT172. A search in the Dali protein structure comparison server showed that these WD domain conformations are unique in solved or Alphafold-predicted human structure databases.

**Figure 3.**
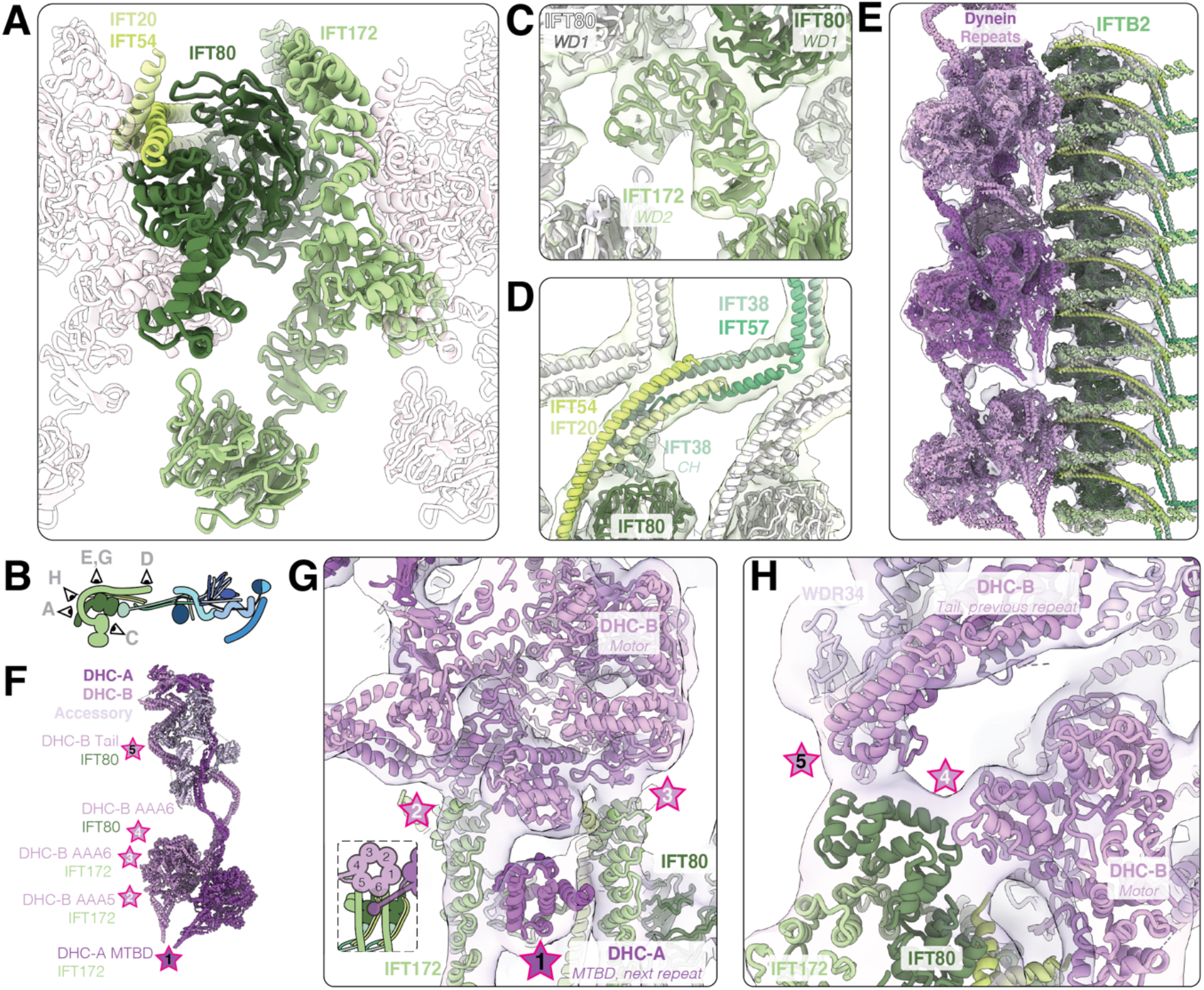
The interaction between IFTB2 and dynein-2. **A** − IFT80 (dark green) forms the core of the IFTB2 complex. It is surrounded by IFT172 (olive green) and the IFT54/20 (lime green, pale green) coiled-coil. Adjacent repeats shown in silhouette **B** − Cartoon representation of IFTB depicting the position of the views in the other panels **C** − The second WD domain of IFT172 (olive green) does not close into a ring, and bridges two IFT80 subunits (dark green from the same complex, white in the neighbour). **D –** In the center of the complex, IFT54/20 (lime/pale green) and IFT57/38 (turquoise/mint green) coiled coils stack on top of each other, stabilizing a kink in IFT57/38 to point the subsequent coiled coils towards IFTB1 **E** − The flexibly refined dynein models (purple, pink) docked into the 16Å dynein density, along with the IFTB2 model. **F** − Cartoon representation of cytoplasmic dynein-2 refined into our density, with the points that contact IFTB2, and the protein they interact with, highlighted with stars. **G** − Top view of the train, showing the first three contact points between dynein and IFTB2. **H** − The two remaining contact points between dynein and the edge of IFTB2, at the C-terminus of IFT80.

From our structure we see that IFT80 is at the center of the IFTB2 subcomplex, with much of its surface covered by protein interactions (**Figure 3A/B)**. The IFT80 WD domains are sandwiched between the WD and TPR domains of two neighbouring copies of IFT172 (**Figure 3A/C)**. Previous work suggested that IFT80 homodimerizes in the initial TPR region^15^, but it is monomeric in our average. Instead, IFT80-TPR wraps around the N-terminal TPR motifs of IFT172 from the neighbouring repeat. IFT172 contains an extended TPR domain that is not reinforced through the formation of a superhelical twist like IFT88/70, meaning that it is likely to be more conformationally flexible. The remaining IFT172-TPR region wraps around the edge of IFTB2 and runs towards the center of the train, forming the roof of the complex (**Figure 2A)**. In summary, IFT80 organises both the core architecture of the IFTB2 complex as well as forming an extended lateral interface capable of stabilizing flexible domains upon polymerization.

### IFT57-CH prevents IFT172-WD1 from interacting with membranes

The IFT172 WD domains were previously shown to bind to and remodel membranes in vitro, suggesting that IFT172 may play a role in membrane trafficking given its structural homology to COPI/II protein family members^25^. However, membrane binding was mutually exclusive with an interaction between IFT57-CH and IFT172-WD. We wanted to see if this interaction is present in active anterograde trains. In our structure, IFT172-WD1 protrudes like a foot from the periphery of IFTB2 and was resolved to lower resolution due to its flexibility. However, masked refinement of this region shows a clear bulge in the density that can be explained by IFT57-CH binding to IFT172-WD1 (**Figure S7E)**. This interaction is made possible in our model by the long unstructured linker between IFT57-CH and the C-terminal coiled coil region that interacts with IFT38 (**Figure S5A)**. This therefore suggests that IFT57-CH helps remove IFT172 from its putative membrane-trafficking phase and makes it available for incorporation into assembling trains.

### The coiled coils in IFTB are in a compressed conformation

Similar to IFT81/74 of IFTB1, a segmented coiled coil in IFTB2 formed by IFT57/38 is folded into a compressed conformation through lateral interactions with neighbouring repeats. IFT57/38 is anchored to IFTB2 through the IFT38-CH/IFT80 interaction (**Figure S6G)**. This is supplemented by the formation of a short four-helix bundle with IFT54/20, which is a single continuous coiled coil that bridges the gap in IFT80-WD2 and runs down to the center of the train (**Figure 3A, S7F**). The helical bundle forms lateral interactions with IFT57/38 in the neighbouring repeat, stabilizing a kink between segments to point it towards the IFTB1 subcomplex (**Figure 3D)**. This is a second right angle corner between IFT57/38 segments stabilized by the neighbouring repeat, after the contact with IFT88 in IFTB1 (**Figure S6H)**. We previously showed that retrograde trains have a much longer repeat than anterograde trains (∼45nm vs. 11.5/6nm), despite being made of the same constituents^9^. We hypothesize that the compressed coiled coils in anterograde trains can be utilised during remodelling by extending into elongated conformations while mantaining intra-complex interactions.

### IFTB cargo binding regions are found on the exterior of the complex

The main role of anterograde IFT is to deliver structural and signalling cargoes from the cell body to the cilium. Biochemical studies have identified several interactions between these cargoes and individual IFT proteins, which we are now able to pinpoint to specific locations of the train. The axonemal outer and inner dynein arms are linked through their specific adaptors to IFT46 and IFT56 respectively^4,26,27,28,29^. These large structural cargoes will therefore be docked on the peripheral surface of IFTB1 (**Figure S8A)**. Furthermore, the N-terminus of IFT70 is located on the same patch of IFTB1, and is thought to recruit a variety of membrane proteins in humans and *Chlamydomonas* ^30,31^ This region of the train, on the opposite side to the dynein-2 binding site, presents the largest open surface of IFTB and was observed to contain heterogeneous extra densities in raw electron tomograms^9^. Therefore, we would anticipate that other large structural cargoes (e.g. radial spokes, dynein-nexin regulatory complex) would be engaged in similar interactions with the same IFT proteins.

Soluble tubulin is an IFT cargo thought to be recruited by a tubulin-binding module composed of IFT81-CH and the basic N-terminus of IFT74 ^14,32^. In our structure, the residues in IFT81-CH important for tubulin binding lie in a narrow gap between coils that prevents an interaction (**Figure S8B)**. Alternatively, IFT81-CH could bind to tubulin in the same way as the highly structurally conserved CH domain of kinetochore protein Ndc80^33^ (**Figure S8C**). However, this would lead to strong steric clashes with IFT81/74 in neighbouring repeats (**Figure S8D)**. This leaves the possibility that the IFT81/74 module binds to the acidic and unstructured C-termini of tubulin, although this would be an unusual way for a CH domain to bind tubulin.

### The cytoplasmic dynein-2 binding sites are only formed upon IFTB polymerisation

The retrograde IFT motor dynein-2 is also transported as a cargo of anterograde trains to the tip of cilia, where it is used to transport retrograde trains back to the cell body. Previously, we showed that autoinhibited dynein-2 complexes dock onto IFTB in a regular repeat, on the edge of what we now determine to be IFTB2^9^. We wanted to understand the molecular basis for this recruitment, however the dynein density was averaged out of our initial structure since its repeat is three times that of IFTB. To address this, we used unsupervised 3D classification to identify a sub-class of particles where all the dyneins are in the same register. We then performed local refinements on this sub-class to obtain an improved 16.6Å final map of dynein-2, and flexibly fit the single particle structure of human dynein-2^34^ into it **(Figure 3E, S7G-I)**.

The dynein dimer consists of two heavy chains (DHC-A/B) that are split into an N-terminal tail domain and a C-terminal AAA+ motor domain^34^. The tail is used for dimerization and recruitment of accessory chains, and the motor domain generates force and binds to microtubules through a microtubule-binding domain (MTBD). The dynein-2 structure underwent few changes during flexible fitting, except for a shift in the C-terminus of the DHC-A into the density (**Figure S7J)**. This left an extra density linking the tail of DHC-A to the motor of DHC-B, which we assign to be the Tctex1 dimer that was weakly resolved in the single particle structure^34,35^. The formation of the Tctex1 bridge could help stabilize the autoinhibited form of dynein in the train.

Dynein-2 binds to IFTB2 at five contact points (**Figure 3F-H**). The first is a composite surface between two IFTB2 complexes that is only formed upon polymerisation. Here, the MTBD of DHC-A sits in a trench formed between the TPR domains of two neighbouring IFT172 subunits, with IFT80-WD2 and IFT54/20 forming the base. This interaction could be mediated by a negatively charged patch on the side of IFT80-WD2, mimicking the interaction between the MTBD and the negatively charged surface of microtubules (**Figure L/M)**. Two more contacts are made by the motor domain of DHC-B bridging the same two IFT172 subunits through the AAA5 and AAA6 domains. The other side of the DHC-B AAA6 domain makes an additional contact with the C-terminal TPR motifs of IFT80 (**Figure 3F-H)**. Finally, the tail of DHC-B from the adjacent dynein repeat contacts the same region of the IFT80 TPRs. These contacts could be supplemented by additional, unstructured contacts, like the reported interaction between the disordered N-terminus of IFT54 and dynein^36^.

Therefore, we find that dynein-2 is only able to bind to IFTB2 in the context of an assembled anterograde train. Its binding site includes the TPR domain of IFT172, which is stabilized in trains but is likely to be flexible in solution based on the Alphafold2 ensemble confidence predictions. This, combined with the MTBD binding site that sits on the boundary between IFTB repeats, means that dynein will only be able to form weak interactions with unpolymerized IFTB. This provides a level of regulation to prevent dynein-2 binding to individual IFTB components before train assembly. Furthermore, it supports the theory that dynein-2 adopts the open conformation ready for retrograde transport directly upon anterograde train disassembly, as a result of the loss of binding sites that stabilise the autoinhibited conformation^34^.

### The IFTA polymer is continuously interconnected

The IFTA complex sits between the IFTB complex and the membrane (**Figure 1B**). In anterograde trains it is responsible for transport of some membrane cargoes. In retrograde trains, IFTA is the complex that binds to active dynein-2, bringing IFTB and retrograde specific cargo back to the cell body. IFTA is made up of five structural proteins (IFT144/140/139/122/121) and one disordered protein (IFT43). IFT144, IFT140, IFT122 and IFT121 all have tandem N-terminal WD domains followed by extended TPR domains (**Figure S5A**). IFT139 consists solely of TPR repeats, which were predicted by Alphafold2 to form a superhelical spiral. However, how these proteins are organised into the IFTA complex, and how the complexes assemble into polymers could not be resolved in previous studies.

The resolution of our IFTA reconstruction was limited to 18.6Å, potentially making subunit placement difficult. However, the Alphafold2 models of each of the four WD-containing IFTA proteins showed unique combinations of angles between the two WD domains and the position of the first TPR repeat (**Figure S9A-D)**. This allowed us to unambiguously place the WD domains in our map, and fit the C-terminal TPR domains into the connected continuous tubular densities. Finally, we identified a spiral density corresponding to IFT139 to complete our model **(Figure S9E/F, Supplemental movie 3)**. We also see an extra density at lower thresholds bridging the gap between the WD domains of IFT144 and IFT140 (**Figure S9G)**. IFT43, which is predicted to be mostly unstructured, is the only IFTA protein that we do not include in our model. However, since IFT43 is thought to interact with two proteins (IFT121 and IFT139^16,18^) that we show are at the other end of the complex, it is unlikely that this density corresponds to IFT43. Therefore, the density belongs to another, unidentified protein.

Our model demonstrates that IFTA is an intricately interconnected complex. IFT144-WD defines one end of the IFTA complex (**Figure 4A-C)**, and projects out towards the membrane. The IFT140-WD domains are nearby, and the N-terminal TPR motifs of IFT144 and IFT140 have a long interface running along the edge of the complex (**Figure 4B)**. Surprisingly, the end of IFT140-TPR runs into the neighbouring repeat, where it interacts with the TPR domain of the adjacent copy of IFT144 (**Figure S9H/I)**. This interaction supports the end of IFT144-TPR, which acts as the base on which IFT140-WD and IFT121-WD sit.

**Figure 4.**
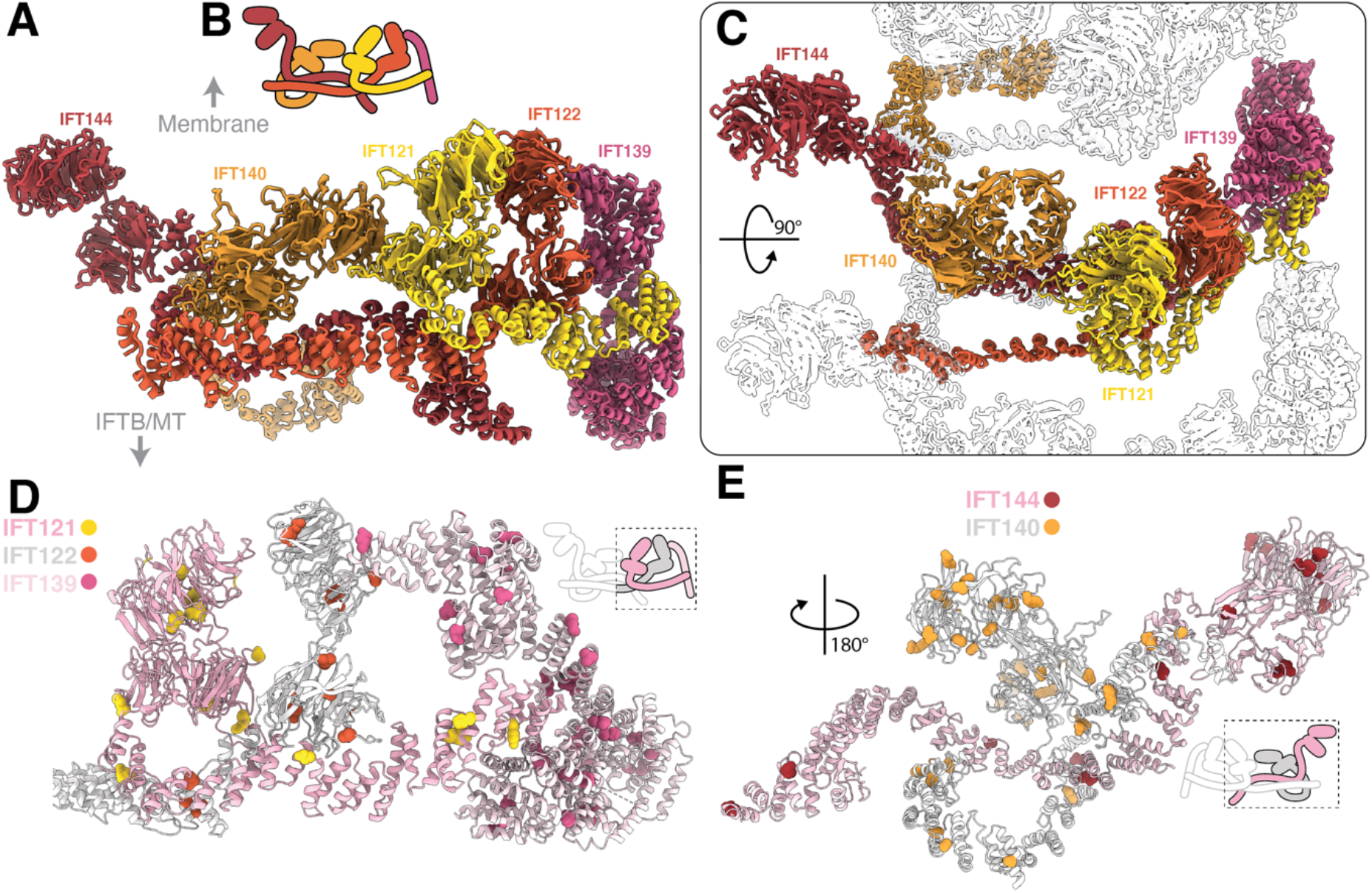
IFTA presents its four WD domains to the membrane. **A** − The IFTA model viewed in profile, as if looking down the train **B** − Cartoon representation of IFTA shown with a side view as in A **C** − Top view of the IFTA model, with neighbouring repeats shown as silhouettes. IFT140 (orange) and IFT122 (light red) both form part of the adjacent complex. **D** − We mapped human point mutations in IFTA proteins that are linked to ciliopathies to conserved residues in C. reinhardtii. Here, IFT121, IFT122 and IFT139 are shown, with most point mutations (shown as sphere representation) mapping to the WD domains or to interfaces between TPR domains. **E** − A second view, showing the point mutations present in IFT144 and IFT140

IFT122, IFT121 and IFT139 form three pillars at the other end of IFTA. The IFT122 and IFT121 WD domains are stacked together directly below the membrane. IFT121-TPR runs through this region to form a platform for IFT122-WD binding and slots into the IFT139 superhelix. (**Figure 4A)**. Finally, IFT122-TPR projects out of the columns towards the adjacent repeat, where it interacts with for IFT144-WD (**Figure S9J/K)**. This unusual arrangement means that IFT140 and IFT122 are responsible for both lateral interactions, and the fundamental structural organisation of the neighbouring repeat. Thus, the most striking feature of IFTA is the interconnectivity between adjacent repeats.

### IFTA mutations are clustered around interfaces

There are over 100 point mutations in IFTA proteins associated with ciliopathy phenotypes in the Human Gene Mutation Database^37^. To understand how these mutations disrupt normal function, we mapped their positions to the homologous residues in *Chlamydomonas* (**Figure 4D/E, Supplementary data 1)**. Many of the mutations can be mapped to the outer surfaces of the WD domains. Since these regions all directly face the membrane, mutations here could have a deleterious effect on membrane recognition or cargo binding. In IFT144 and IFT140, many of the WD domain mutations correspond to the regions that interact with the unidentified extra density (**Figure S9G)**. This suggests that this extra density could be an IFTA cargo or cargo adaptor.

In the TPR domains, almost all the mutations are found at the interfaces with other IFTA proteins (**Figure 4D/E)**. This includes interactions between IFT144 and IFT140 belonging to neighbouring repeats (**Figure 4E)**. These mutations are therefore likely to result in destabilization of the complex, due to disruption of complex formation or polymerization. IFT139 is an exception because it contains mutations throughout its structure. It forms an external surface of the complex, meaning mutations are likely to disrupt interactions with cargo or IFTB (discussed below) rather than complex formation.

### IFTA and IFTB are flexibly tethered

A major remaining question is how IFTA and IFTB can stably bind to each other given their periodicity mismatch. In our overall IFTA and IFTB averages, the mismatch meant that one complex was blurred out in the average of the other (**Figure 5A-C)**. By using masked 3D classification of the region corresponding to IFTA in our IFTB averages, we obtained classes where IFTA is resolved in different registers relative to the IFTB (**Figure S10A)**. In these classes, we see two new densities bridging IFTA and IFTB (**Figure 5D/E**).

**Figure 5.**
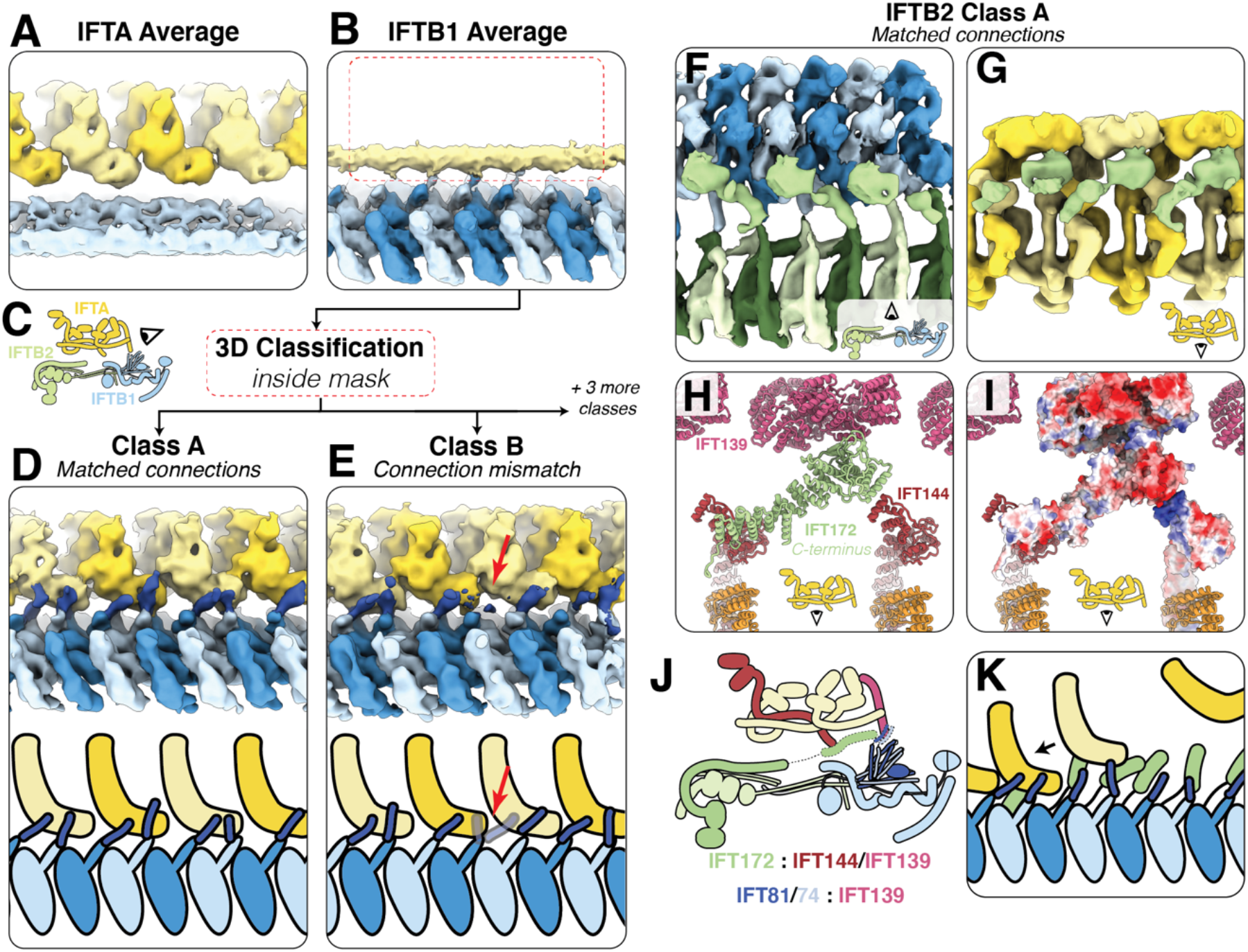
IFTA and IFTB are connected at two points. **A** − The 21Å IFTA average covering three repeats, unmasked to show that IFTB (light blue) is averaged out with respect to IFTA (alternating yellow) due to peridocity mismatch **B** − The IFTB1 average filtered to 12Å and unmasked, to show that IFTA (yellow) is averaged out with respect to IFTB1 (alternating blue) due to periodicity mismatch. Red box indicates the location of the mask used for subclassification to generate the classes in D/E **C** − Cartoon depicting the view in A, B, D and E **D** − After classification of the IFTA region in the IFTB1 average, we find classes where IFTA (alternating yellow) and IFTB (alternating blue) are in sync. We see a new density (dark blue) linking IFTB to IFTA, which we designate as CC5 of IFT81/74. Bottom, cartoon representation of the density. **E** − A second class shows how the IFT81/74 connections (dark blue) adapt to the periodicity mismatch between IFTA (alternating yellow) and IFTB (alternating blue), by switching register with respect to IFTA at the red arrow. Bottom, cartoon representation of the density. **F** − A top view of Class A from classification of the IFTA region in the IFTB2 average (cartoon view shown inset). IFTB1 (alternating light/dark blue) and IFTB2 (alternating light/dark green) are joined by a new, unmodelled density corresponding to the C-terminus of IFT172 (lime green). **G** − The same class as **F**, rotated 180° to view the same IFT172 density (lime green) interacting with IFTA (alternating yellow). Cartoon view inset. **H** − The same view as **G**, showing the Alphafold2 IFT172 C-terminus model (lime green) docked into the density along with our IFTA model. IFT172 bridges the gap between IFT144 and IFT139. **I** − The same view as **H**, with IFT172, IFT144 and IFT139 shown with surface charge depiction. The negatively charged IFT172 C-terminus can make favourable ionic interactions with the positively charged IFT144 C-terminus **J** − Cartoon representation of the overall anterograde train structure, showing the two points of connection (dotted outlines).

The first bridge is between IFT139 in IFTA and IFT81/74 in IFTB1 (**Figure 5D)**. Each IFTB1 repeat projects a tubular density corresponding in length and location to the unmodelled fifth coiled coil segment of IFT81/74. Two copies of IFT81/74 bind to one IFT139, although there are transition zones where the periodicity mismatch means two adjacent repeats are competing for the same IFT139 binding site (**Figure 5E**). Here, there is a switch in register in the subsequent repeats, made possible by the conformational flexibility between IFT81/74 coiled coil segments. IFT139 has a strongly negatively charged surface and IFT81/74-CC5 is positively charged, making a favourable ionic interaction possible (**Figure S10B/C)**. This interaction is also consistent with the mutations in IFT139 we find in this region (**Figure 4D**), which could affect IFT81/74 binding.

The second IFTA/IFTB bridge is visible in classes obtained from our IFTB2 average. We see an extension of the IFT172 density running along the roof of IFTB2 in alternate repeats (**Figure 5F/G)**. This density reaches up to the IFTA complex and docks between the C-terminus of IFT144 and the inner face of IFT139. We assign this density to be the C-terminal TPR domain of IFT172, which is also unmodelled in our overall model. Like IFT81/74-CC5, this domain is linked to the modelled region by a flexible linker, allowing it to interact with IFTA in a range of registers. The C-terminus of IFT172 contains a strongly acidic patch capable of binding to a basic patch on IFT144 (**Figure 5H/I)**. We only see the extra density extending from alternate IFTB repeats, reflecting the fact that this binding site is only present once per IFTA repeat.

Together, we show that anterograde trains overcome the periodicity mismatch between IFTA and IFTB using flexible tethers from IFTB that are in a stoichiometric excess to IFTA. This mode of interaction provides several advantages for the function of anterograde trains. Firstly, it suggests that IFTA is recruited in a search-and-capture mechanism, where nascent IFTB polymers can sample a large space through these tentacle-like tethers (**Figure 5K)**. This then aids IFTA polymerization by creating a higher local concentration of IFTA to promote their lateral interaction into polymers. In principle, this could mean IFTA could only polymerise with the help of IFTB, thus preventing IFTA multimerisation away from the basal body. Finally, a flexible interaction allows IFTA and IFTB to maintain their connection while withstanding the mechanical stresses present in actively beating cilia

## Discussion

The key outstanding question is how the structure we show here remodels into the conformationally distinct retrograde train. We recently showed that anterograde to retrograde train conversion in *Chlamydomonas* can be induced by mechanical blockage of IFT at arbitrary positions along the length of the cilium^38^. This indicates that anterograde to retrograde remodelling does not require specialized machineries of the ciliary tip. In *Chlamydomonas*, the constituents from one anterograde train appear to split into two or three retrograde trains, with IFTA and IFTB complexes remaining associated during anterograde to retrograde conversion^39^. Together, this supports a model in which conversion occurs through conformational changes pre-built into the anterograde train. This could be through the compressed or spring-like coiled coils such as IFT81/74 or IFT57/38. Alternatively, TPR and other alpha-solenoid domain proteins have previously been shown to behave as molecular springs^40–42^. Many of the TPR domains in our structure underwent curved-to-straight conformation changes to fit the relaxed Alphafold2 predictions into our density (**Figure S5B**), indicating that they could be a source of molecular strain. This strain could then be released at the tip, potentially triggered by the loss of tethering to the microtubule, resulting in a relaxation into the retrograde conformation.

Consistent with this model, retrograde trains that are mechanically blocked in the cilium never convert back into anterograde trains^38^. This suggests that the compressed structures that we see in anterograde trains require an external packaging mechanism during train assembly. Interestingly, in subtomogram averages of anterograde trains assembling at the basal body, an unknown extra density is observed beneath IFTB1 that is absent in the mature train^3^. This unknown component could therefore be what loads the molecular springs in the anterograde train. However, to fully understand how this, and how train conversion occurs, more structural information of the retrograde train is required.

## Methods

### Cell culture

*C. reinhardtii* wild-type (CC625) cells and CC625 cells with glycocalyx proteins FMG1A and FMG1B deleted by CRISPR (produced for and described in a manuscript in preparation) were cultured in aerated Tris-acetate-phosphate (TAP) media at 24°C with a 12/12 hour night/dark cycle for at least two days before use.

### Grid preparation

Quantifoil R3.5/1 Au200 grids were plasma cleaned for 10s with a 80:20 oxygen:hydrogen mix (Solarus II Model 955, Gatan). 4uL cells were added to grid, followed by 1uL 10nm colloidal gold fiducial solution (in PBS, BBI Solutions). Following 30s incubation at 22°C at 95% humidity, the grid was back-blotted and immediately plunge frozen in liquid ethane at −182°C (Leica Automatic Plunge Freezer EM GP2).

### Cryo-electron Tomography data acquisition

Cryo-ET data were acquired on a Thermo Scientific Titan Krios G4 transmission electron microscope operated at 300 kV using SerialEM^43^. Raw movie frames were recorded on a Thermo Scientific Falcon 4 direct electron detector using the post-column Thermo Scientific Selectris-X energy filter. Movies were acquired in EER format^44^, with a pixel size of 3.03Å/px, an exposure of 3s and a dose rate of 2.6e^-^/Å^2^/s. Tilt series were collected in 3° increments using a dose-symmetric scheme with two tilts per reversal up to 30°, and then bidirectionally to 60°. For a full tilt series this resulted in an accumulated dose of 104e^-^/Å^2^. Tilt series were acquired between −2.5 and −4.5um defocus.

### Tomogram reconstruction

Tilt series reconstruction was performed using a developmental update of the TOMOMAN pipeline^45^, which organises tomographic data while feeding it into different pre-processing programs. Motion correction was performed using the MotionCor2 implementation in Relion3.1^46^, with EER data split into 40 fractions. Bad tilts were then removed after manual inspection, followed by dose weighting (Imod^47^) and CTF estimation (CTFFIND4^48^). Manual fiducial alignment and CTF-corrected tomogram reconstruction at bin4 was then performed in Etomo^47^. The bin4 tomograms were then deconvolved for visualisation with the tom_deconv filter^49^.

### Particle Picking

Anterograde IFT trains were identified in deconvolved bin4 tomograms according to features identified previously^9^. Picking was performed using the 3DMOD slicer^47^, with IFTB and IFTA picked separately. For each IFTB and IFTA filament, a open contour model was picked along the length. Points were picked along this contour at 4/2nm distances for IFTA/B respectively (representing a ∼3x oversampling in each case) using Tom Toolbox scripts^50^.

### Subtomogram averaging

We used STOPGAP^51^ to find initial orientations before transferring data to Relion for high resolution refinements. However, we found that because IFTB looks similar with 180° rotation around the long axis (phi angle in STOPGAP) the initial angles were split roughly 50/50 with the right and wrong phi angle. We therefore analysed each train individually and determined a rough phi angle manually. In STOPGAP, we extracted particles from the unfiltered bin4 tomograms (70/50px box sizes for IFTB/IFTA) and performed alignments using a cone search with 32° phi search in 8° increments.

The particles and orientations from STOPGAP were converted to Relion star format, and subtomograms and 3D CTF particles were extracted in Warp^52^.

For IFTB, six different collection sessions were incrementally added to the average (**Figure S2)**. Each group was refined separately in STOPGAP, with the STOPGAP average of the first group used as the initial reference for 3D refinement in Reolion 3.1^46^. Initial refinements used a solvent mask consisting of the entire IFTB complex for four repeats. We performed a local 3D refinement with 3.7° initial angular sampling/step, and 4/1 pixels search/step. The resulting refinement was used as the input for a round of image warp grid refinement in M^53^. The refined subtomograms were re-extracted and the 3D refinement was repeated, resulting in a significantly improved average. This refinement was then used as the input for 3D classification into two classes, using the same solvent mask and keeping the alignments fixed. The particles from the good class were then used for separate masked refinements of IFTB1 and IFTB2, which proceeded independently but with the same input particles. For IFTB1 we found that reducing the length of the mask to 2 repeats resulted in the best averages, but IFTB2 was best at 4 repeats. Both subcomplexes reached Nyquist resolution, so IFTB1 was reextracted eventually to bin 1 (3.03Å/pix) and IFTB2 to bin 1.5 (4.04Å/pix). We obtained the highest resolution reconstructions after performing image warp and ctf refinement on the IFTB1 reconstruction in M. We used the resulting parameters to reextract both IFTB1 and IFTB2 particles for a final round of 3D refinement (1.7°, 3/1). Resolution was determined with the 0.143 threshold (**Figure S3A/B)**. Masked refinement of the ends of IFTB1 and IFTB2 resolved these regions more clearly, although still at lower overall resolution compared to the core masks (**Figure S2C)**. To obtain an average of dynein, we created a solvent mask based on our previous low-resolution IFTB/dynein average and rescaled it to 4.04Å/px (**Figure S2D)**. We performed 3D classification on our IFTB2 average into 6 classes without refinement (**Figure S2A**), finding three classes with dynein in three registers. We selected one class and performed local refinement.

For IFTA, the six collection session groups were combined directly after STOPGAP into a local refinement in Relion using a mask with three repeats (**Figure S4)**. We did not perform image warp refinement in M for IFTA, as it resulted in a worse average compared to when the refinements from IFTB1 were used. However, we found that after the first refinement in Relion, we saw a strong improvement by applying the median Phi angle for each train to every particle in the same train (coordinate smoothing). This pulls particles that have strayed back to the consensus angle for the train. The smoothed coordinates were then locally refined in Relion again, and this refinement was used for masked 3D classification without alignments. The good class reextracted at bin2 (6.06Å/px) and locally refined with a selection of masks (one repeat, three repeats, left side and right side (**Figure S4B-E)**) to generate maps that best show individual features within the complex and also connections between adjacent complexes.

### Model building

A number of crystal structures were available for IFTB components, but we used Alphafold2 structural predictions for all components because the crystal structures were either from different species or only contained fragments of the protein. Structure predictions were run as monomers or multimers using a local install of Alphafold version 2.1.1 ^54^. Alphafold2 predictions had no major differences to the solved crystal structures. All IFTA proteins were folded as monomers. For IFTB, IFT172 and IFT56 were the only proteins folded as monomers. In IFTB1, the complexes folded as multimers were IFT88-52-70, IFT70-52-46^11^ and IFT81-74^13^. For IFT70, the best fit of the density was achieved by splitting the model in two, with the IFT88-52-70 prediction contributing the C-terminus and the IFT70-52-46 contributing the C-terminus. IFT52 was split at the same place as IFT70. In IFTB2, we folded IFT80-57-38 and IFT54-20 as multimers ^10,15^.

Once we had these starting models, the position of most of the IFTB proteins in the density was straightforward. IFT172, IFT88/70/52, IFT81/74, IFT80 all contained strong structural motifs that us to position the original Alphafold2 models unambiguously. This left the two coiled coil densities in IFTB2 to fill. Based on the known interaction between IFT80 and IFT38-CH, we pinpointed the IFT38-CH domain to the density bound to the face of IFT80-WD1. From here, the length of the three IFT57/38 coiled coil segments exactly matched the coiled coil density that reaches across from IFTB2 to IFTB1. Finally, the length of IFT54/20 matched the coiled coil density running down the side of IFT80, consistent with the unstructured IFT54 N-terminus interacting with cytoplasmic dynein-2.

For IFTA, the four proteins with WD domains each contain unique conformations regarding the angle between the tandem WD domains, and between the second WD domain and the start of the TPR. This allowed us to place each of the four WD domains into the density unambiguously. We recognized that the proteins could not adopt reasonable conformations to fit into one repeat as defined in our previous cryo-ET structure. However, we could identify continuous density between adjacent repeats in the average of three consecutive IFTA repeats. The IFT139 TPR superhelix was obviously identifiable at the edge of the complex, but was split into two rigid bodies at a loop in the middle of the protein to best fit the density.

Once we had positioned the models in the density, we manually edited them to best fit the density. In IFTB1, in regions where individual alpha helices were resolved (IFT88, IFT70, IFT81/74, IFT57/38) this involved conventional secondary structural real-space refinement in Coot ^55^. In IFTB2, the IFT54/20 coiled coil needed to be curved slightly to fit into the density. The C-terminal TPR domains of IFT172 curved out of the density. To counter this, we split the region into rigid bodies defined by loops where the Alphafold2 prediction had lower confidence. We then fit the rigid bodies up to the point where the density became too weak, leaving roughly one third of IFT172 unmodelled (Figure S5A). We used the same approach for the TPR domains in IFTA. For IFT140, IFT122 and IFT121 we did not model the flexible TPR regions at the very C-termini. This is because they were predicted to be only loosely tethered to the remaining TPR regions, but in each case there is empty density left in the average for them to occupy.

Once we had manually assembled the models into the density, we used NAMDinator^56^, an automated molecular dynamics flexible fitting (MDFF) pipeline, to refine to models into our density. We used default parameters, and started with the individual assemblies described above. Different models were then combined to form the IFTB1/2 and IFTA complexes and refined, and then combined again to create lateral repeats to ensure lateral did not clash. Map and model visualization was performed in ChimeraX ^57^. Human point mutations were obtained from the Human Gene Mutation database ^37^.

## Acknowledgements

We would like to thank P. Swuec and S. Sorrentino from the Human Technopole electron microscopy facility, and C. Fernandez and P. Margara for IT and HPC support; P. Erdmann and S. Khavnekar for providing the TOMOMAN and STOPGAP implementations; D. Diener, A. Vanninni, F. Coscia for comments on the manuscript; A. Nievergelt for CRISPR modified cell lines.

## Author contributions

S.E.L. prepared the samples, acquired cryo-ET data, performed image processing, refined and analysed the data and wrote the manuscript. H.E.F. performed Alphafold2 structural predictions. G.P. designed the experiments, analysed the data and wrote the manuscript.

**Figure S1.**
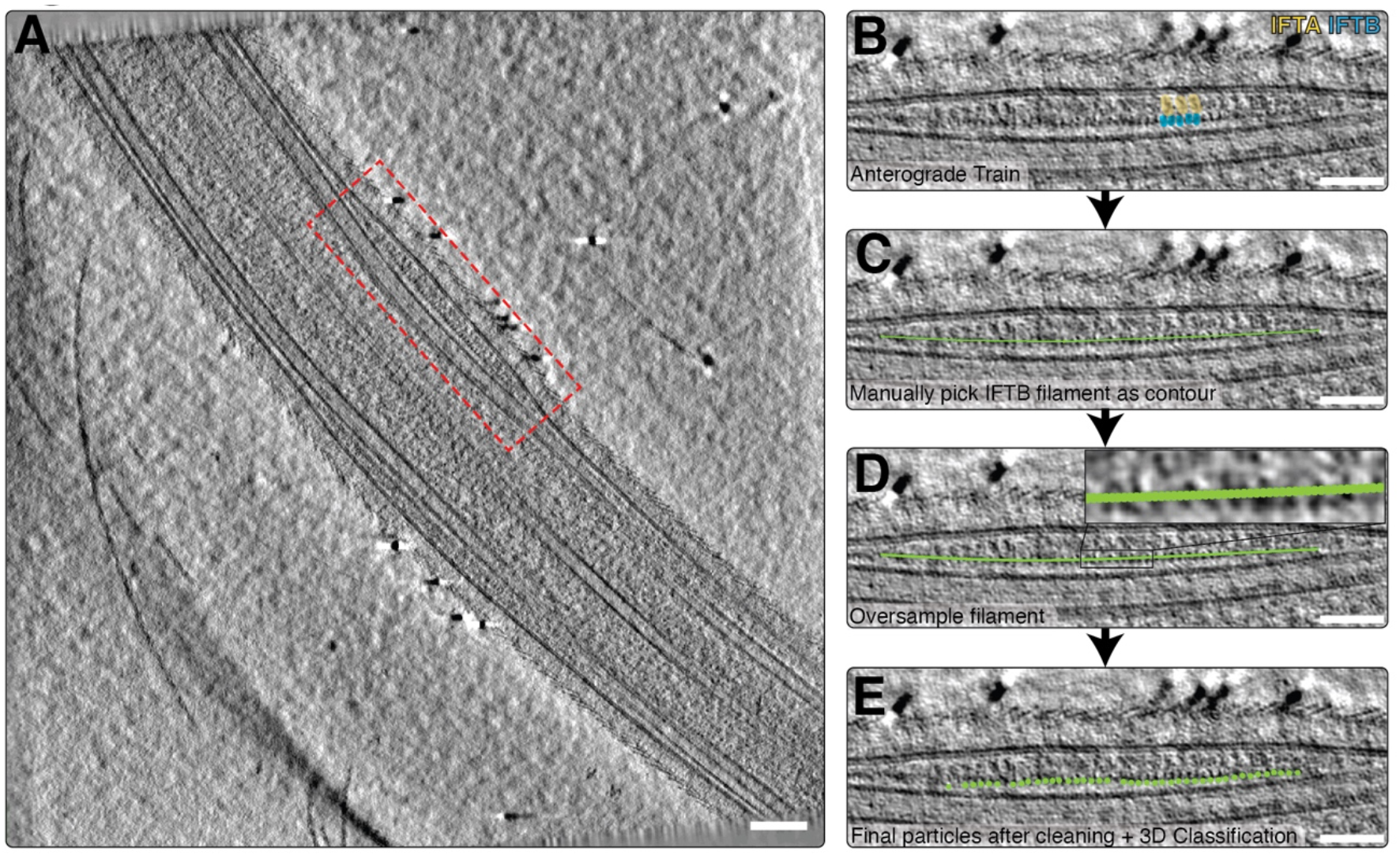
Identification of anterograde IFT trains in cryo-electron tomograms. **A** − A slice through a tomogram of a C. reinhardtii cilium, showing a bulge in the membrane in the middle corresponding to an anterograde IFT train (red box). Scale bar=100nm **B –** Close up view of the train in A, with IFTA (yellow) and IFTB (blue) repeats annotated. Scale bar=50nm **C** − After identification, we manually picked trains in IMOD as a contour running through the center of the complex. IFTB picking is shown here, and IFTA, visible above the IFTB contour, was picked in a separate model. Scale bar=50nm **D** − The contour was converted into subtomogram coordinates with oversampling to ensure no particles were missed. Scale bar=50nm **E** − Here, the final refined coordinates are shown on the train. The particles have undergone proximity cleaning compared to the oversampling in D, as well as 3D classification to remove bad particles. Scale bar=50nm

**Figure S2.**
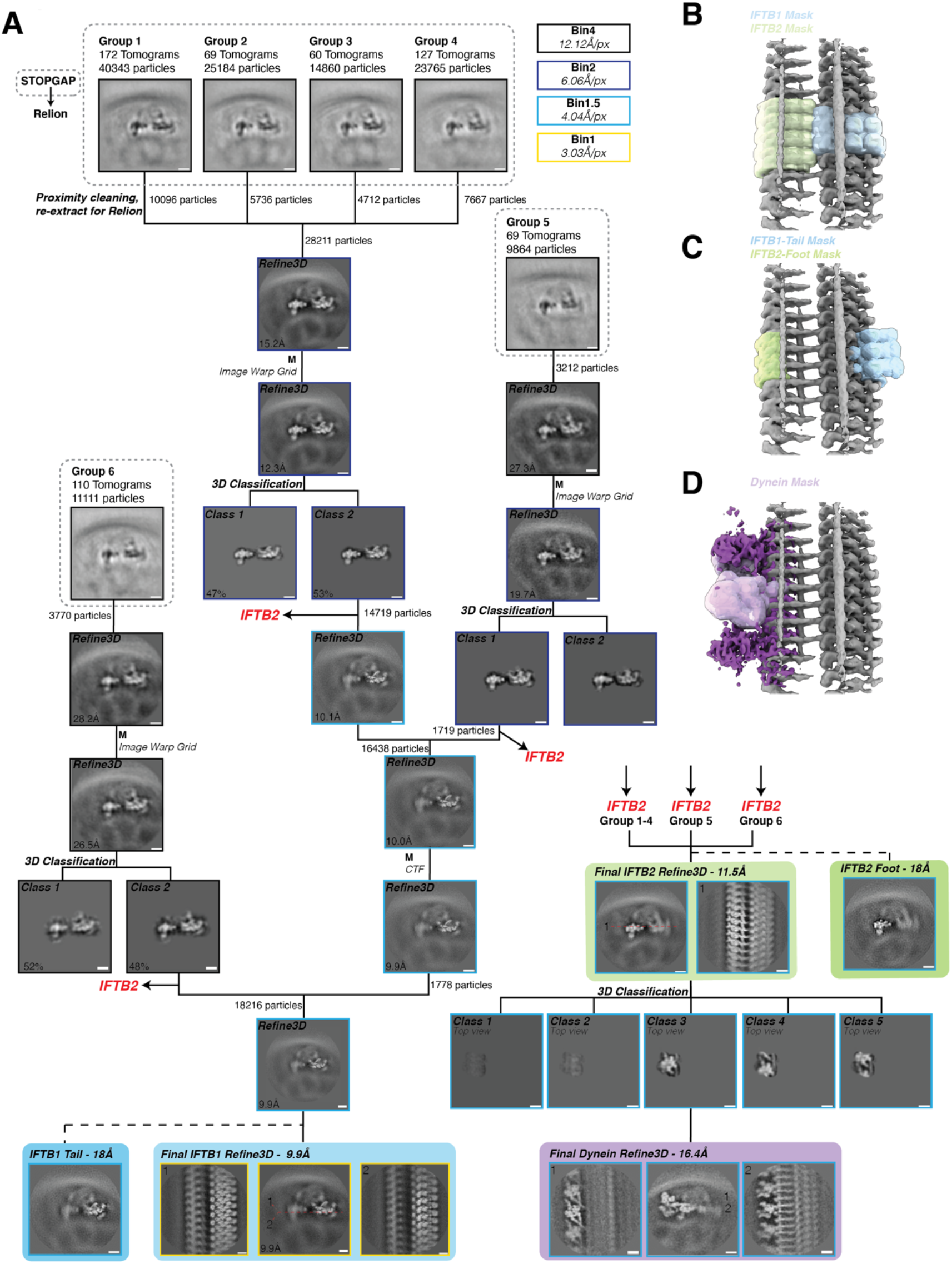
Processing diagram for IFTB subtomogram averaging. **A –** Workflow depicting the steps involved in averaging the IFTB1 and IFTB2 complexes. Processing started in STOPGAP (areas in dotted black line) before proceeding to Relion. The level of binning at each stage is indicated by the outline of the box (colour code top right).All scale bars=10nm **B –** The solvent masks used to refine IFTB1 (blue) and IFTB2 (green) separately from each other **C –** The solvent masks used to refine the extremities of the IFTB1 and IFTB2 complexes, which are poorly resolved when using the masks in B **D –** The solvent mask used to classify and refine dynein from IFTB2.

**Figure S3.**
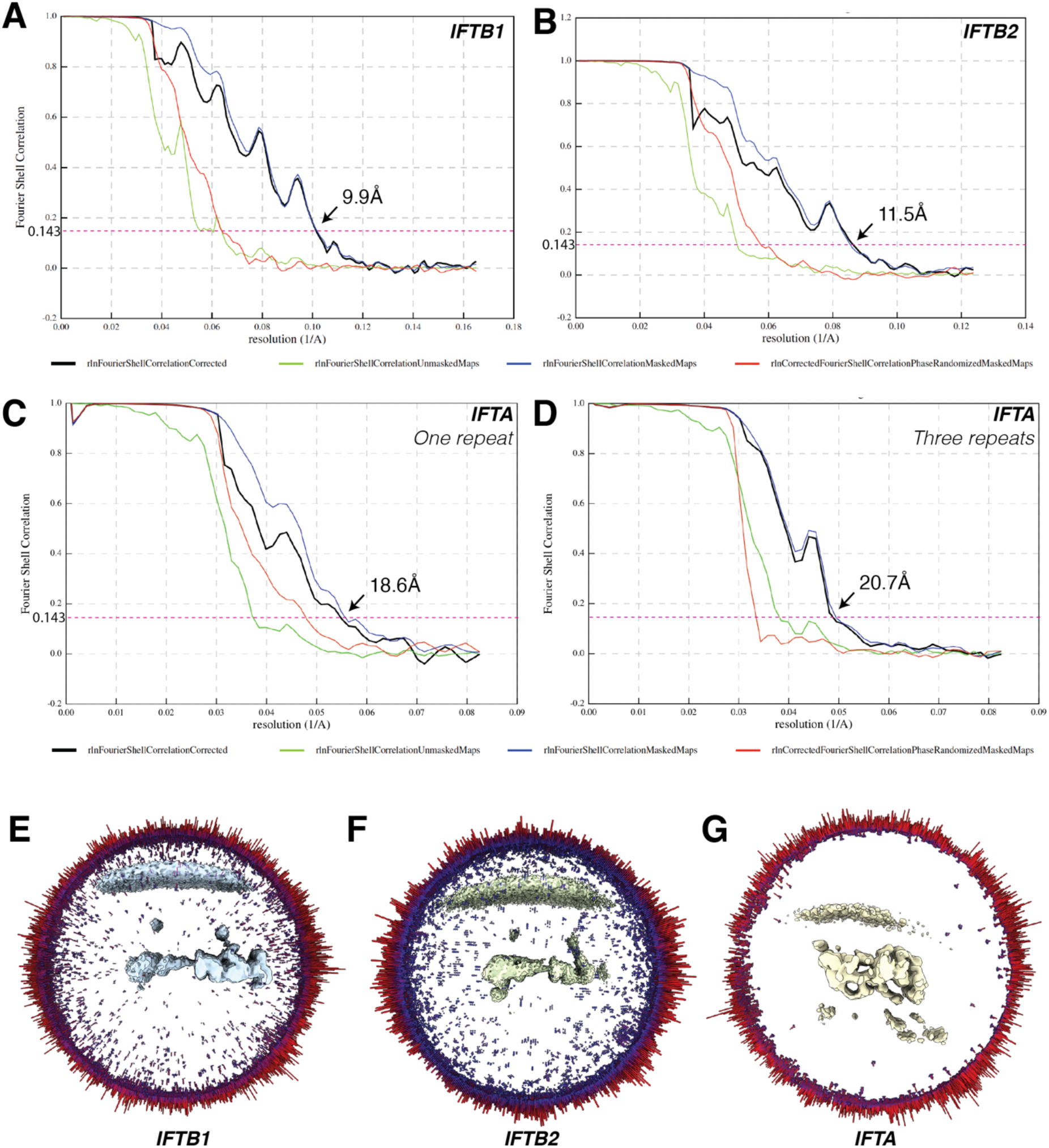
Descriptions of IFTB and IFTA map quality. **A –**Fourier Shell Coefficient (FSC) curve of the IFTB1 average, as a measure of map resolution **B –** FSC curve of the IFTB2 average **C –** FSC curve of the IFTA average, refined using a mask containing one repeat **D –** FSC curve of the IFTA average, refined using a mask containing three repeats **E –** Angular distribution of particles contributing to the IFTB1 average **F –** Angular distribution of particles contributing to the IFTB2 average **G –** Angular distribution of particles contributing to the IFTA average (one repeat)

**Figure S4.**
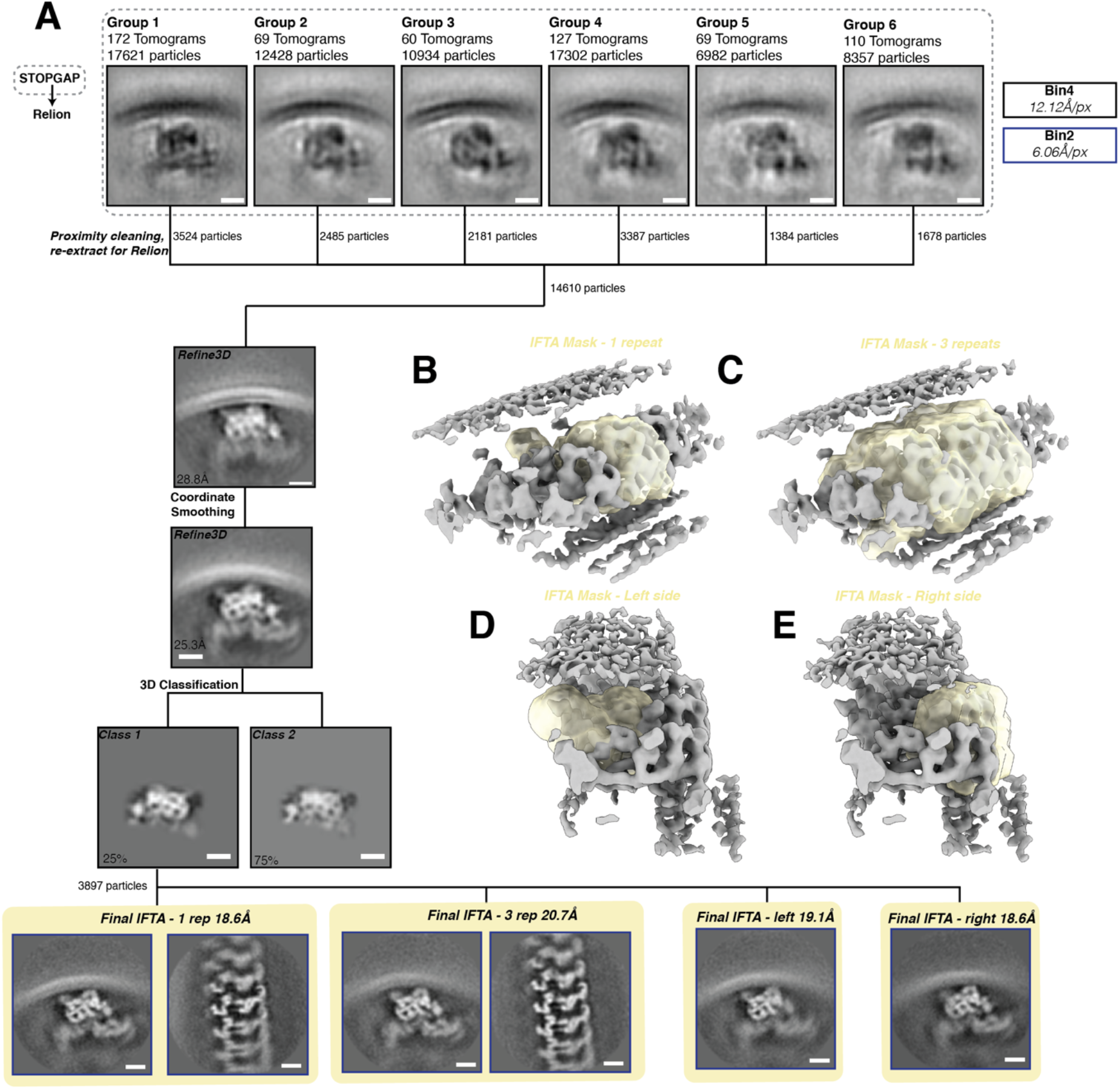
Processing diagram for IFTA subtomogram averaging. **A** − Workflow depicting the steps involved in averaging the IFTA complex. Processing started in STOPGAP (areas in dotted black line) before proceeding to Relion. The level of binning at each stage is indicated by the outline of the box (colour code top right). All scale bars=10nm **B –** The solvent mask used to refine IFTA, containg one repeat **C –** The solvent mask used to refine IFTA, containg three repeats **D –** The solvent mask used to refine IFTA, consisting of the left side of one repeat of the complex **E** − The solvent mask used to refine IFTA, consisting of the right side of one repeat of the complex

**Figure S5.**
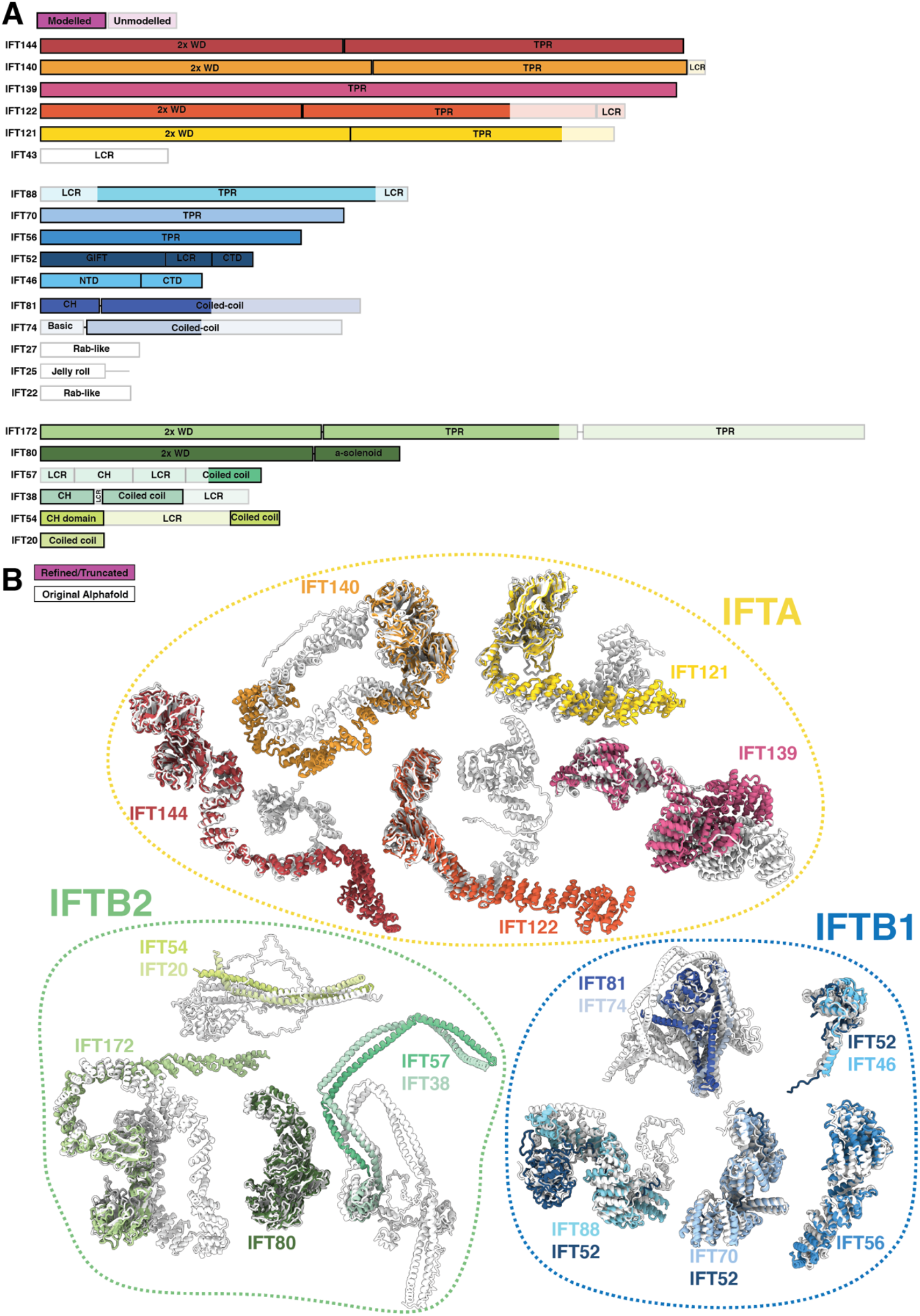
Building a model of IFT using Alphafold2 predictions. **A** − Domain organization of all IFT constituents. Lighter shading indicates regions that were flexible and unmodelled in our structure. WD = WD40 repeat domain, TPR = Tetratricopeptide repeat domain, CH = Calponin homology domain, LCR = low-complexity (disordered) region. **B** − The original, unmodified alphafold structures (white) overlaid with the final refined models in our new structure (colours). Refined models have had flexible regions deleted.

**Figure S6.**
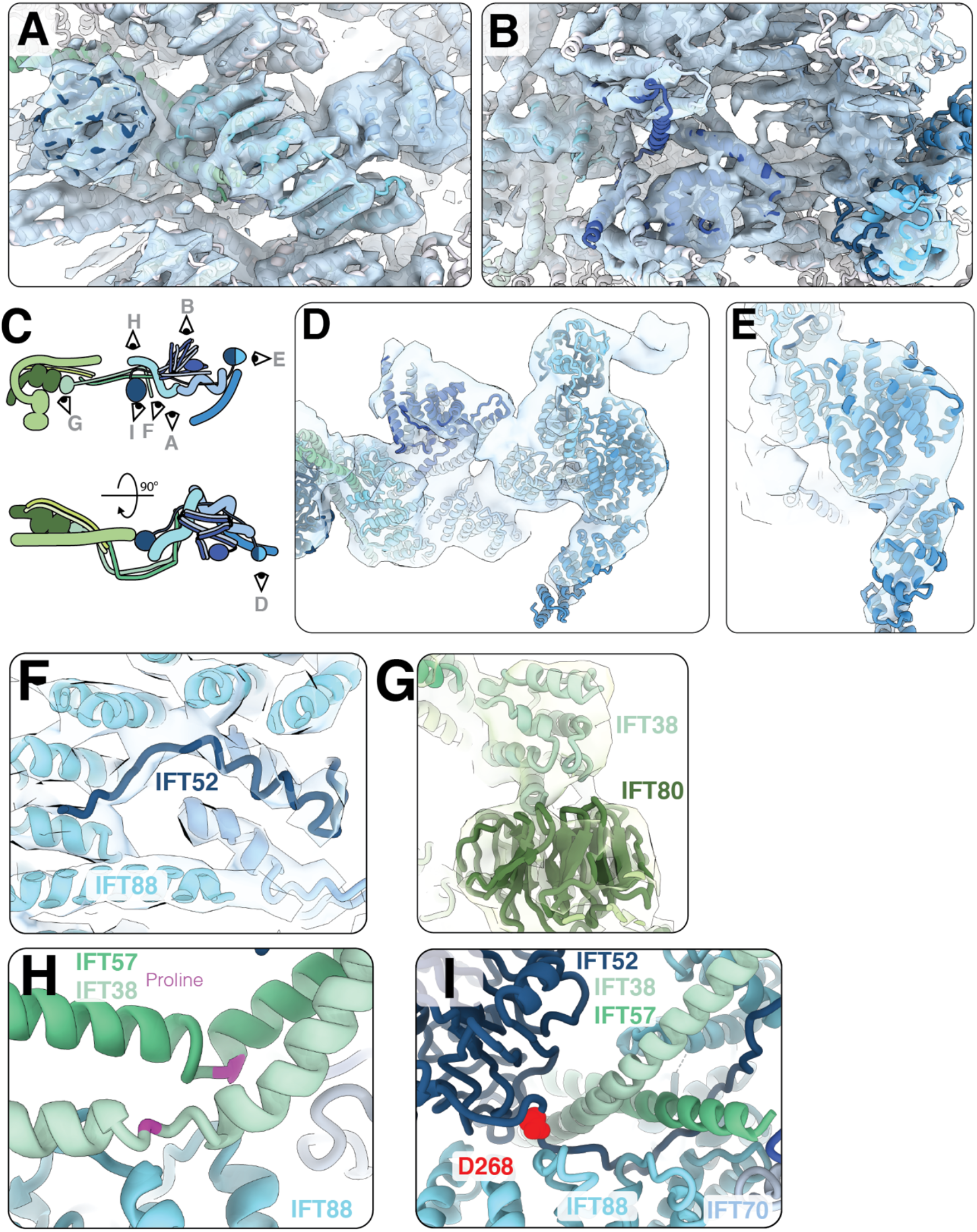
Building a model of IFTB1. **A –** A view of the IFTB1 model docked into its density from the bottom (see E) **B –** A view of the IFTB1 model docked into its density from the top (see E) **C –** Cartoon representation of IFTB showing the views in A-D. **D –** A side view of the “tail” of IFTB1 docked into the masked tail refinement (Figure S2A) map lowpass filtered to 18Å. The region containing IFT56 was more flexible in the high-resolution average shown in A/B, but is more clearly resolved here. **E –** A close up view of IFT56 in the masked tail refinement map, showing that the twist in the TPR helix is visible **F –** Density for the central unstructured domain of IFT52 (dark blue) is visible in the central pore of IFT88 (cyan), showing that the Alphafold2 prediction agrees with our experimental data. **G –** The N-terminal CH domain of IFT37 (light green) docks to the exterior face of the first WD domain of IFT80 (dark green) in IFTB2. **H –** A proline residue (magenta) creates a kink in each of the IFT57/38 (dark/light green) helices near the contact to the first IFT88. **I –** The position of D268 in IFT52 highlighted in red, at the interface between IFTB1 and IFTB2. D268 inC. reinhardtii corresponds to the D259H mutation in humans ^22^.

**Figure S7.**
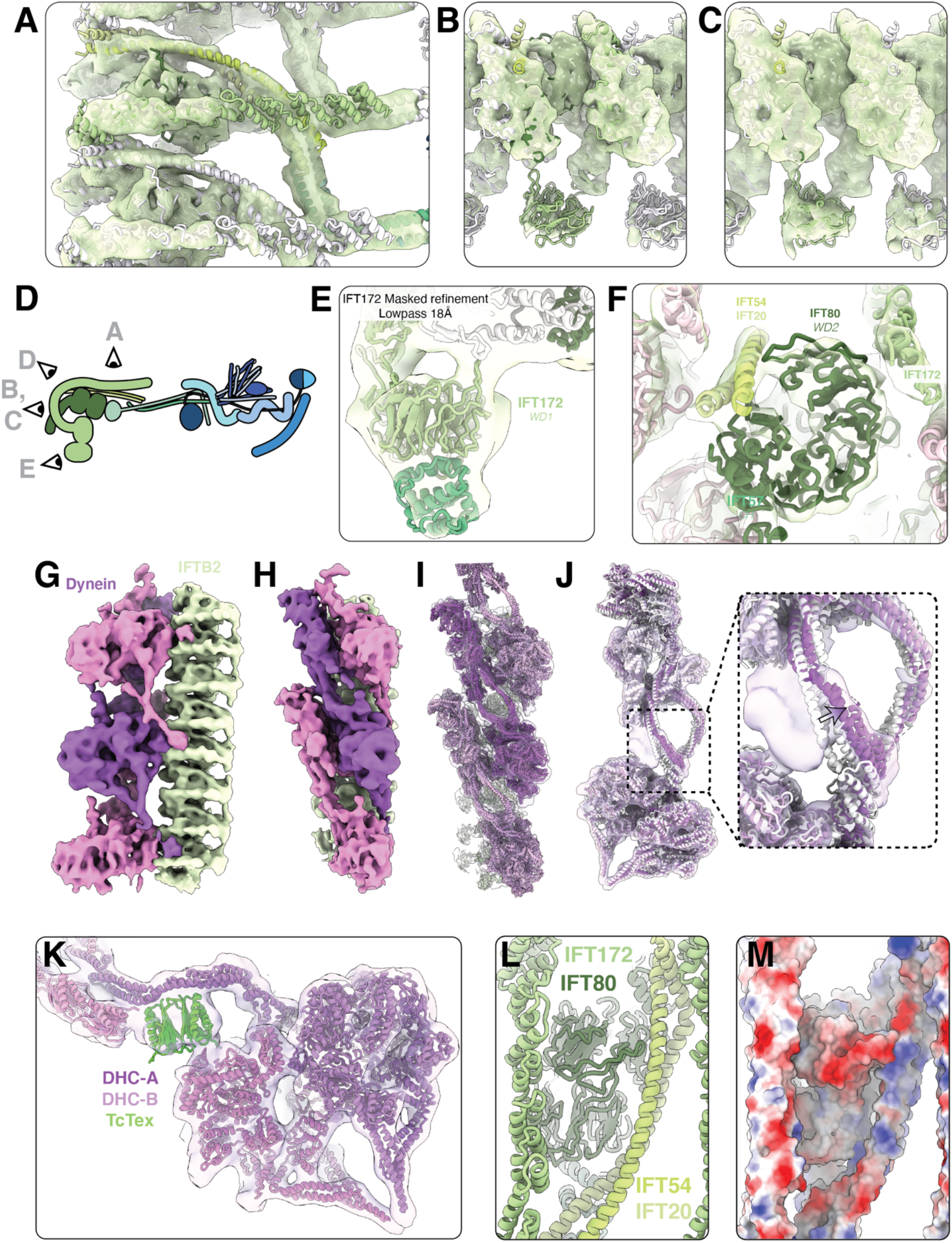
Building a model of the IFTB2 complex and its interaction partner dynein-2. **A –** A top view of the IFTB2 subtomogram average density with the IFTB2 model docked in. **B –** A view of the end of the IFTB2 subtomogram average density with the IFTB2 model docked in. **C –** The same view as B, but at a lower threshold to demonstrate that IFT172-WD1 is represented in the density but at lower resolution than the rest of the complex due to flexibility. **D –** Cartoon depicting the views of IFTB in the other panels **E –** The IFT172-WD1 domain folded as a multimer with the CH domain of IFT57 forming a complex that is represented in the density of the IFT172 masked refinement map. **F –** The IFT54/20 (lime/pale green) bridge the gap in the IFT80-WD2 ring. **G –** Coloured density of Figure 3D, showing our newly refined dynein average. Dynein repeats are alternating pink/purple, IFTB2 is green **H –** Side view of F **I –** Same view as G, with density made translucent and the models docked in. **J –** The density in our new dynein average cropped out around the original dynein model (white) shows that the heavy chain undergoes a rearrangement in our newly refined model (purple), leaving an unmodelled density (inset). **K –** The unmodelled density likely corresponds to a Tctex1 dimer (green), linking the motor domains to the tail. **L –** A view of the top surface of IFTB2, corresponding to the site where the dynein MTBD binds. **M –** The same view with surface charge representations shown, highlighting a positively charged patch where dynein binds.

**Figure S8.**
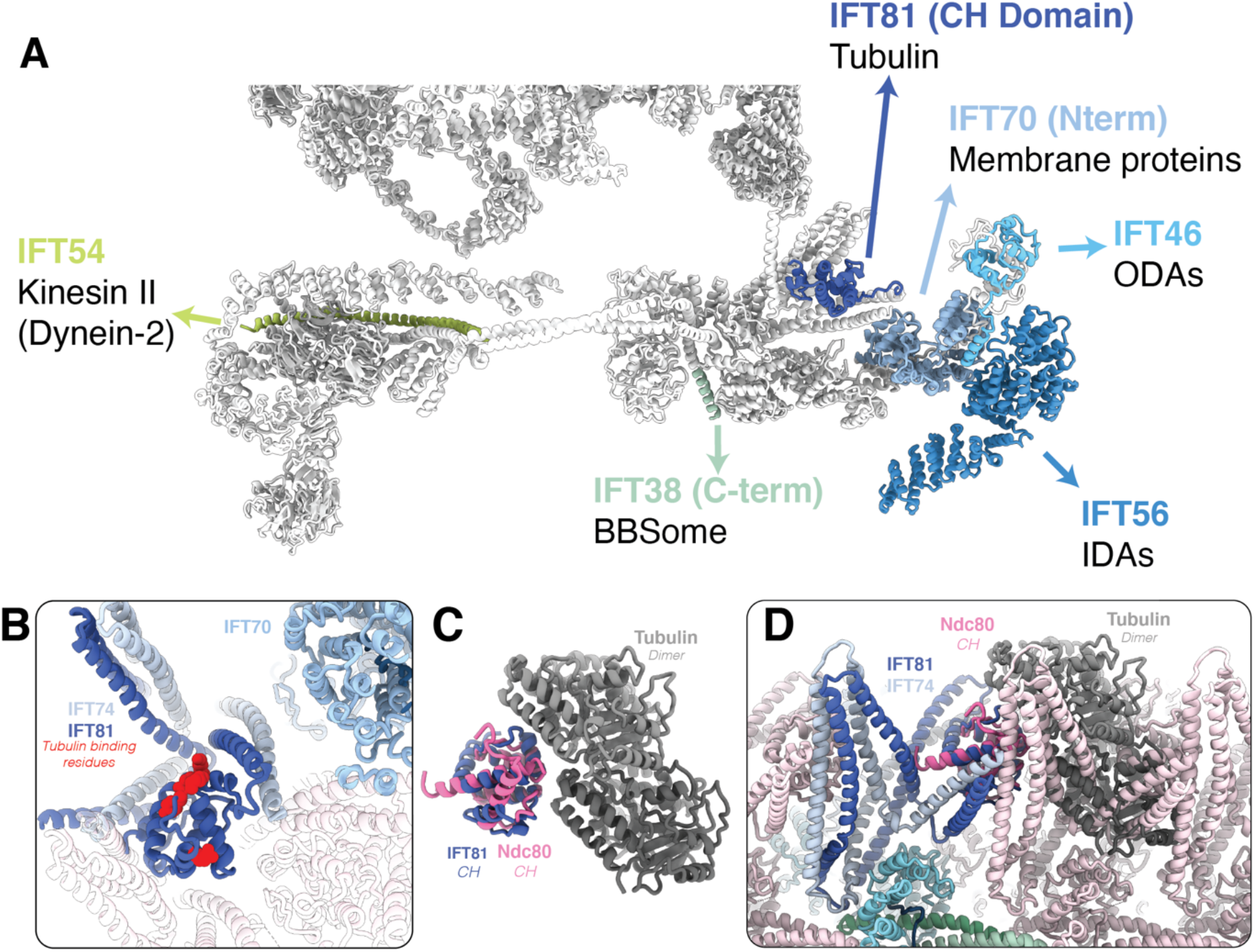
Cargo interactions in anterograde IFT trains. **A** − The IFTA and IFTB models are displayed in grey, with regions of IFTB previously linked biochemically to cargo transport labelled coloured. The large structural cargo interactions mostly occur at the edge of IFTB1. IFT54 is thought to recruit kinesin II to anterograde trains, but this is not visible in our structure, probably due to flexibility. **B** − The CH domain of IFT81 (navy blue), with positive residues thought to be important for tubulin binding shown in red. Only a narrow space exists between the coiled coil domains of IFT81/74 nearby. **C** − Comparison between IFT81 CH domain (navy blue) and the CH domain of Ndc80 (pink) bound to microtubules (grey, PDB 3IZO), indicating strong structural homology between the two CH domains. **D** − The Ndc80:MT complex structure docked with the Ndc80-CH domain aligned to the IFT81-CH domain, simulating a potential interaction with tubulin cargo. Strong steric clashes occur between tubulin and IFT81/74 in the neighbouring repeat.

**Figure S9.**
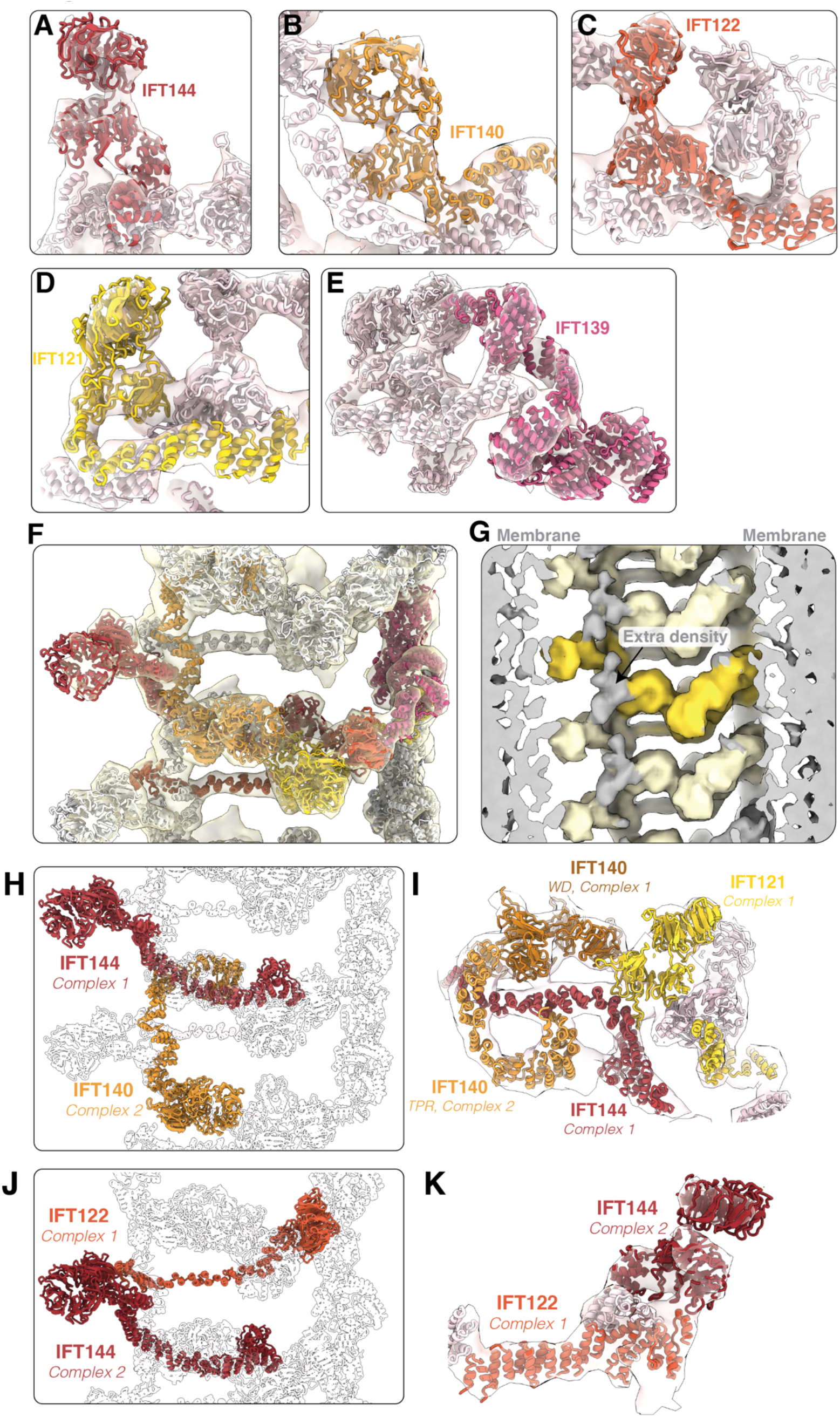
The IFTA polymer is built around four tandem WD domain proteins. **A –** During model building, we located the IFTA subunits based on the conformation of their WD domains. Here, we see the WD domain of IFT144 in the “left masked” IFTA average (Figure S4A). **B –** The WD domains of IFT140 flexibly fit into the “left masked” IFTA average **C –** The WD domains of IFT122 flexibly fit into the “right masked” IFTA average (Figure S4A) **D –** The WD domains of IFT121 flexibly fit into the “right masked” IFTA average **E –** The model for IFT139 flexibly fit into the “right masked” IFTA average **F –** Multiple repeats of the overall IFTA model docked into the “3 repeat” IFTA average (Figure S4A) to show the overall fit of the model into the density. **G –** We lowpass filtered our IFTA 3-repeat average, with regions containing part of our model coloured in yellow (dark yellow highlighting a single repeat). We see an extra density (grey) forming a bridge between the WD domains of IFT144 and IFT140 that is not formed by a protein in our model. **H –** Long distance interconnectivity between IFT144 and IFT140 from neighbouring complexes. The TPR domain of IFT140 (orange) reaches into the neighbouring complex and stabilize its copy of IFT144-TPR (dark red). **I –** Side view of H, with some extra subunits coloured and density shown. The TPR domain of IFT140 from the adjacent repeat (complex 2) stabilizes the conformation of IFT144 (complex 1). The WD domain of IFT140 (dark orange) sits on top of IFT144-TPR (both complex-1), meaning IFT140-TPR from complex 2 determining the conformation of its neighbour. This stabilizes the binding site for IFT121-WD (yellow, complex 1) **J –** Long distance interconnectivity between IFT122 and IFT144 from neighbouring complexes. IFT122-TPR from complex 1 reaches across to form a platform that IFT144-WD from complex 2 sits upon. **K –** Side view of J with density shown.

**Figure S10.**
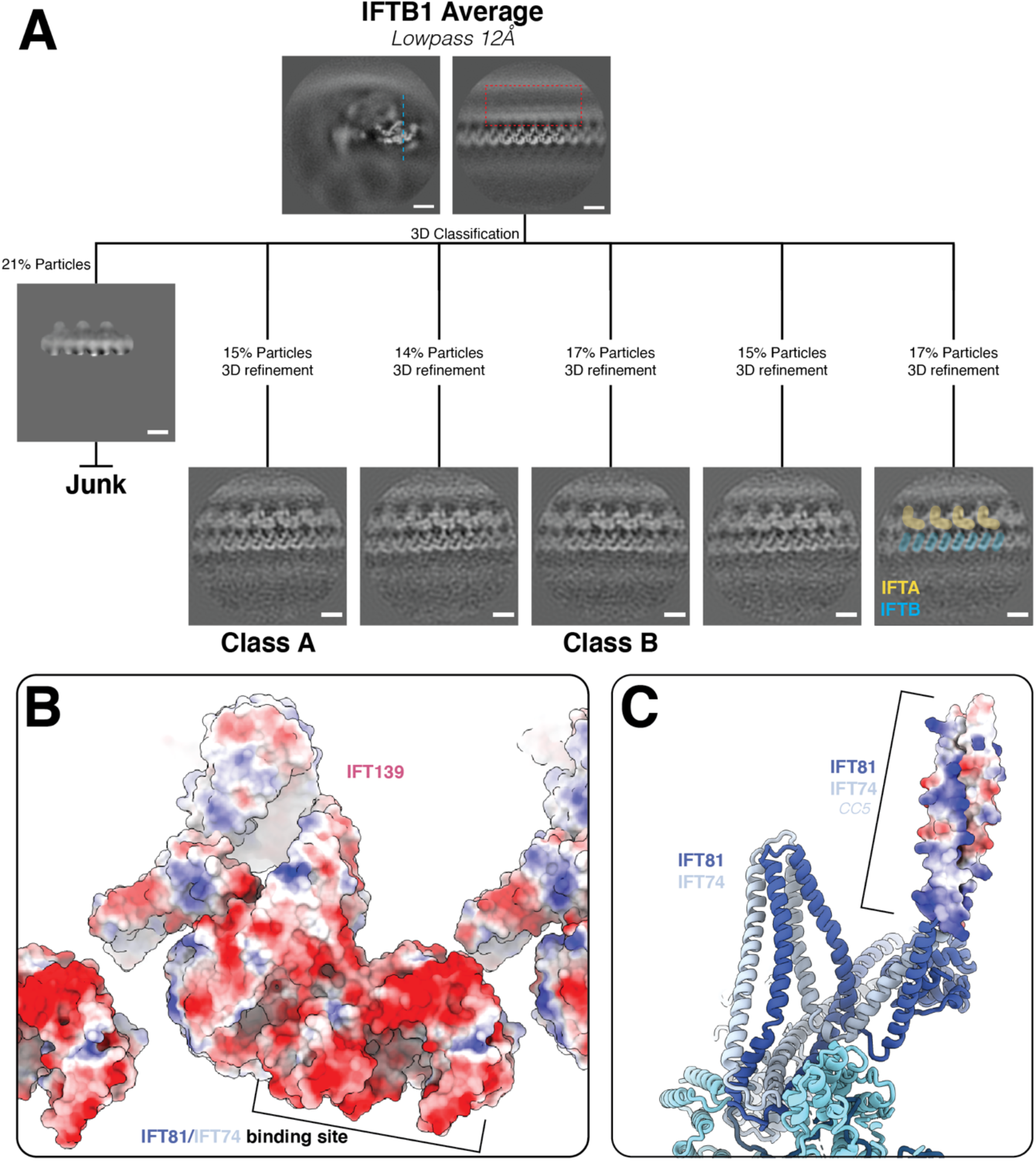
Classification of synchronous IFTA and IFTB averages. **A –** Processing workflow of the classification of the IFTB average to generate the classes in **Figure 5**that show synchronous IFTA and IFTB. Scale bars=10nm **B –** Surface charge representation of IFT139 shows that the IFT81/74 binding site is strongly negatively charged **C –** Surface charge representation of IFT81/74 CC5 shows that it is positively charged, facilitating its interaction with IFT139.

**Supplementary table 1.** Cryo electron tomography data collection parameters

| Data Collection |  |
| --- | --- |
| Microscope | Titan Krios G4 |
| Voltage (kV) | 300 |
| Energy filter | 10-20eV |
| Detector | Falcon 4 |
| Recording mode | EER |
| Pixel size | 3.03Å/px |
| Defocus range (um) | 2.5-4.5 |
| Increments | 3° |
| Tilt range | ±60° |
| Acquisition scheme | Dose-symmetric |
| Accumulated dose (e-/A2) | 100eV |
| Number of tomograms | 600 |

**Supplementary table 2.**
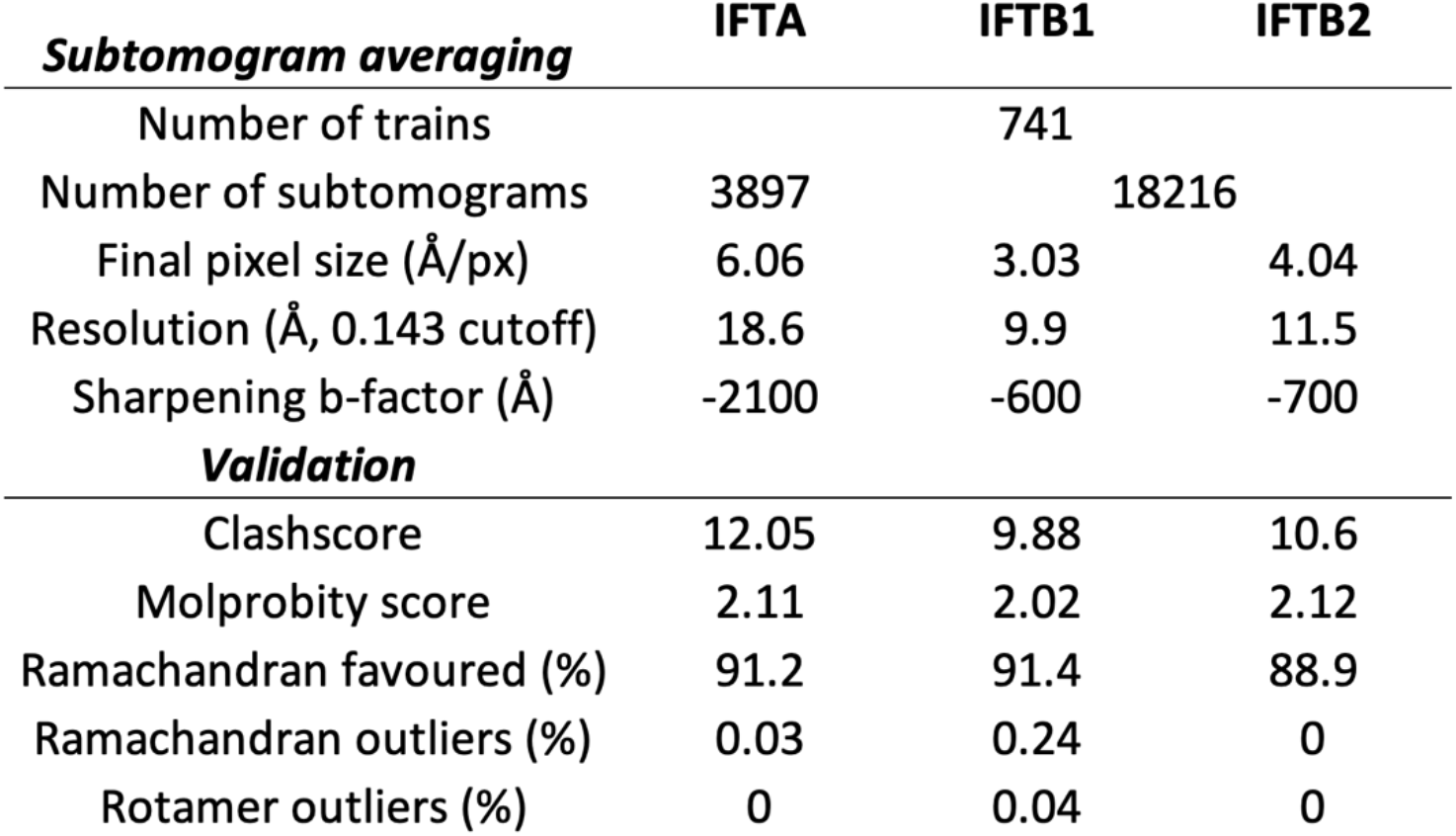
A summary of subtomogram averaging a model refinement validation statistics

**Supplementary Movie 1**

An overview of the IFTB1 complex, showing the fit of the refined model into the density

**Supplementary Movie 2**

An overview of the IFTB2 complex, showing the fit of the refined model into the density

**Supplementary Movie 3**

An overview of the IFTA complex, showing the fit of the refined model into the density

**Supplementary Data 1**

A list of human mutations to IFTA proteins found in the Human Gene Mutation Database, and their corresponding conserved residues in *Chlamydomonas* proteins.

